# The Beta secretase BACE1 drives fibroblasts activation in Systemic Sclerosis through the APP/β-catenin/Notch signalling axis

**DOI:** 10.1101/2022.12.22.521579

**Authors:** Christopher W. Wasson, Enrico De Lorenzis, Eva M. Clavane, Rebecca L. Ross, Kieran A. Walker, Begoña Caballero-Ruiz, Cristina Antinozzi, Rebecca Wells, Gemma Migneco, Jane M. Y. Brown, Natalia A. Riobo-Del Galdo, Luigi Di Luigi, Clive S. McKimmie, Francesco Del Galdo, Paul J. Meakin

## Abstract

The beta-amyloid precursor protein cleaving enzyme 1 (BACE1) is well known for its role in the development of Alzheimer’s disease. Recent publications, including our own, have demonstrated a role for this enzyme in other chronic diseases. The aim of this study was to investigate the role of BACE1 in the autoimmune disease systemic sclerosis (SSc). BACE1 protein levels were elevated in SSc patient skin. Inhibition of BACE1 with small molecule inhibitors or siRNA blocked SSc and fibrotic stimuli mediated fibroblast activation. Furthermore, we show that BACE1 regulation of dermal fibroblast activation is dependent on β-catenin and Notch signalling. The Neurotropic factor BDNF negatively regulates BACE1 expression and activity in dermal fibroblasts. Finally, sera from SSc patients show higher Aβ and lower BDNF levels compared to healthy controls. The ability of BACE1 to regulate SSc fibroblast activation reveals a new therapeutic target in SSc. Several BACE1 inhibitors have been shown to be safe in clinical trials for Alzheimer’s disease and could be repurposed to ameliorate fibrosis progression.

## Introduction

Systemic sclerosis (SSc) is an autoimmune disease that presents with fibrotic involvement of the skin and internal organs including the lungs and heart. The fibrosis is driven by activated fibroblasts (myofibroblasts) characterised by increased expression of α-SMA and secretion of extracellular matrix proteins. Several pathways have been implicated in the activation of fibroblasts in SSc, including the TGF-β, Wnt/β-catenin and Hedgehog pathways (1, 2, 3, 4).

The beta-amyloid precursor protein cleaving enzyme 1 (BACE1) is well known for its role in the development of Alzheimer’s disease via the generation of β-amyloid (Aβ) (5). However, our recent findings have demonstrated a role for this enzyme in other chronic diseases associated with vascular damage, including type-2 diabetes and cardiovascular diseases (6, 7).

Several links exist implicating a potential role for BACE1 in SSc. Vasculopathy is a common feature in SSc pathogenesis (8), developing through the disease. We have shown that BACE1 KO mice are protected from vascular damage in the diet-induced obesity mouse model (9). Further studies showed that inhibition of BACE1 attenuates palmitate-induced ER stress and inflammation in skeletal muscles (10) and ER stress has previously been shown to enhance fibrosis (11). BACE1 can also target the IL-1 receptor 2 (IL-1R2) for cleavage, modulating systemic IL-1 activity through soluble IL1R2 release (12). This is interesting as several IL-1 family members including IL-33 have been implicated in SSc (13). Furthermore, BACE1 expression levels are upregulated by NF-κB through binding of the p65 subunit to the BACE1 promoter (14). Taken together, these data suggest BACE1 levels could be elevated in SSc through inflammatory responses and participate in vasculopathy and fibrosis progression. The aim of this study was to determine if BACE1 plays a role in the pathogenesis of fibrosis in SSc. We show BACE1 protein levels are increased in SSc skin, fibroblasts and a mouse model of fibrosis. Accordingly, the BACE1 products Aβ40 and Aβ42 are higher in SSc patient sera, while the BACE1 negative regulator Brain-derived neurotrophic factor (BDNF) is reduced compared to healthy control. In *in vitro* studies, we show that inhibition of BACE1 activity was sufficient to block expression of pro-fibrotic markers including α-SMA and Collagen type 1 and actin-depending contraction in SSc fibroblasts. Mechanistically, BACE1 promoted fibroblast activation through its ability to activate Notch1 and Wnt3a signalling.

## Materials of Methods

Materials and methods can be found in supplementary file

## Results

### BACE1 expression is increased in SSc patient skin and fibroblasts

To inform the potential role for BACE1 in SSc pathogenesis, we first assessed BACE1 protein expression in healthy (N=5) and SSc (N=5) forearm skin biopsies. BACE1 expression level was much higher in the epidermis than the dermis, both in healthy and SSc patient skin biopsies (Figure 1A). However, dermal BACE1 protein levels were significantly elevated in SSc skin compared to healthy skin. In particular, we observed high levels of BACE1 in fibroblasts and endothelial cells in the SSc dermis (Figure1A inset). The specify of BACE1 staining in the skin samples was confirmed with an isotype control antibody. The high levels of BACE1 staining found in the SSc dermis and epidermis was lost when stained with an isotype control (Supplementary Figure 1)

**Figure 1:**
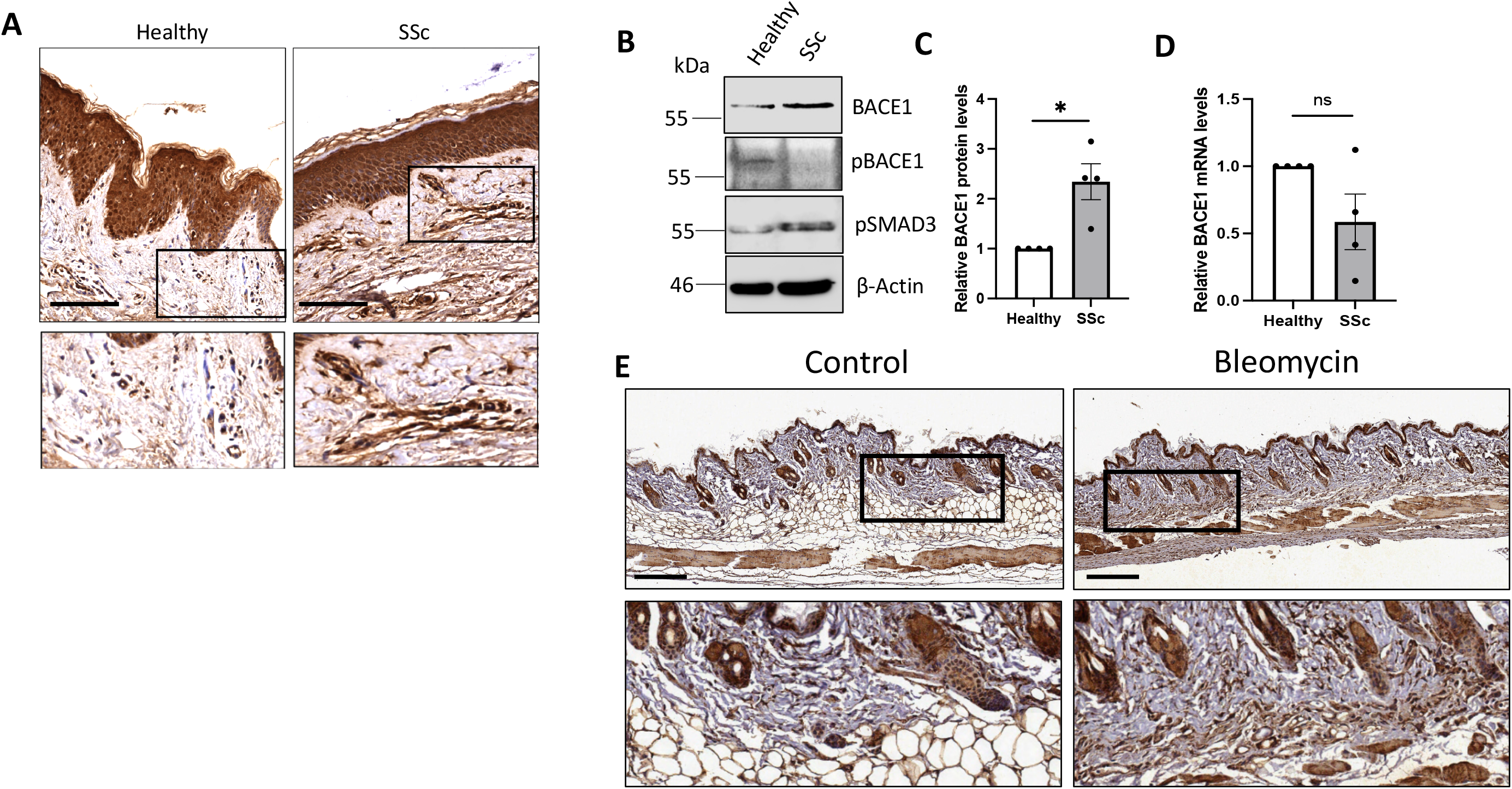
BACE1 levels are increased in SSc patient skin, fibroblasts and in vivo skin fibrosis models. (A) Healthy and SSc patient skin biopsies were stained with an antibody specific to BACE1. Skin sections were counterstained with Haematoxylin. Scale Bars represent 50μM. Protein and RNA were extracted from healthy and early diffuse SSc dermal fibroblasts. (B) BACE1, pBACE1 (Ser498) and pSMAD3 protein levels were analysed by western blot. β-actin was used as a loading control. (C) Densitometry analysis of western blots. (D) BACE1 transcript levels were assessed by qPCR. Graphs represent the mean and standard error. (E) Representative skin sections from vehicle and bleomycin treated mice (N=4 each group) were stained with an antibody specific to BACE1 (Brown). Scale bars represent 200μM. *p<0.05, **p<0.01, ***p<0.001

BACE1 protein levels remained increased in isolated immortalised dermal fibroblasts from early diffuse SSc patients (>2-fold) compared to healthy dermal fibroblasts (Figure1B-C). Similar results were observed between primary healthy and SSc dermal fibroblasts (Supplementary Figure 2A). In addition, we observed high levels of phosphorylated SMAD3 (pSMAD3) in the SSc fibroblasts which confirmed the activated TGF-β phenotype in SSc fibroblasts (Figure1B). Interestingly BACE1 transcript levels did not correlate with protein expression (Figure1D), suggesting that the increase in BACE1 protein levels is regulated at the post-translation level, similar to that seen in Alzheimer’s disease (15). In support of this we found levels of phosphorylated BACE1 at Ser498 were reduced in SSc fibroblasts compared to healthy control and previous studies have shown the phosphorylation of BACE1 can reduce its stability (16)

To assess the role of BACE1 in skin fibrosis establishment *in vivo,* we analysed BACE1 levels in a novel mouse model of SSc by bleomycin treatment of immunodeficient mice after xenotransplantation of human plasmacytoid dendritic cells (pDCs) (17). Subcutaneous injection of bleomycin induced skin fibrosis, loss of subcutaneous adipose tissue and a thickening of the dermis. We observed higher levels of BACE1 in the keratinocytes and endothelial cells in both the control and bleomycin-treated skin (Figure1E); however, BACE1 was specifically upregulated in the dermal fibroblasts in the bleomycin-treated fibrotic skin (Figure1E), similar to that observed in SSc patient skin (Figure1A). This suggests that BACE1 expression is increased in response to induction of skin fibrosis *in vivo*.

### BACE1 plays an important role in SSc and stimuli-induced myofibroblast activation

Next, we set out to determine the role of BACE1 in SSc fibroblast biology using structurally dissimilar small molecule inhibitors against BACE1 (M3 and AZD3839). AZD3839 has completed phase 1 clinical trials for Alzheimer’s disease with a good safety profile (Clinical trial number: NCT01348737). We treated healthy and SSc fibroblasts with these inhibitors and assessed pro-fibrotic gene expression. Both inhibitors led to a reduction in *COL1A2* (Figure2A) and α-SMA (Figure2B) transcript levels in SSc fibroblasts to a level indistinguishable from healthy control fibroblasts. The inhibitors had no effect on fibrotic marker expression in healthy fibroblasts. Similarly α-SMA and CCN2 protein levels were reduced in SSc fibroblasts treated with M3 and AZD3839 (Figure 2C). AZD3839 reduced pro-fibrotic gene expression (*COL1A2, CCN2* and α-SMA) in a dose-dependent manner in SSc fibroblasts (Figure2D-F). M3 had a similar effect in primary SSc and healthy fibroblasts (Supplementary Figure 2B). In the gel contraction assay, AZD3839 (1µM) decreased the contracting ability of SSc fibroblasts (Figure2G). The collagen gels containing SSc fibroblasts weighed less than healthy control gels (Healthy=157mg±41mg, SSc=101mg±14mg). This phenotype was partially ameliorated when the SSc fibroblast gels were incubated with AZD3839 (SSc=119mg±13mg; p=0.0356) but the inhibitor had no significant effect on the weight of collagen gels containing healthy fibroblasts.

**Figure 2:**
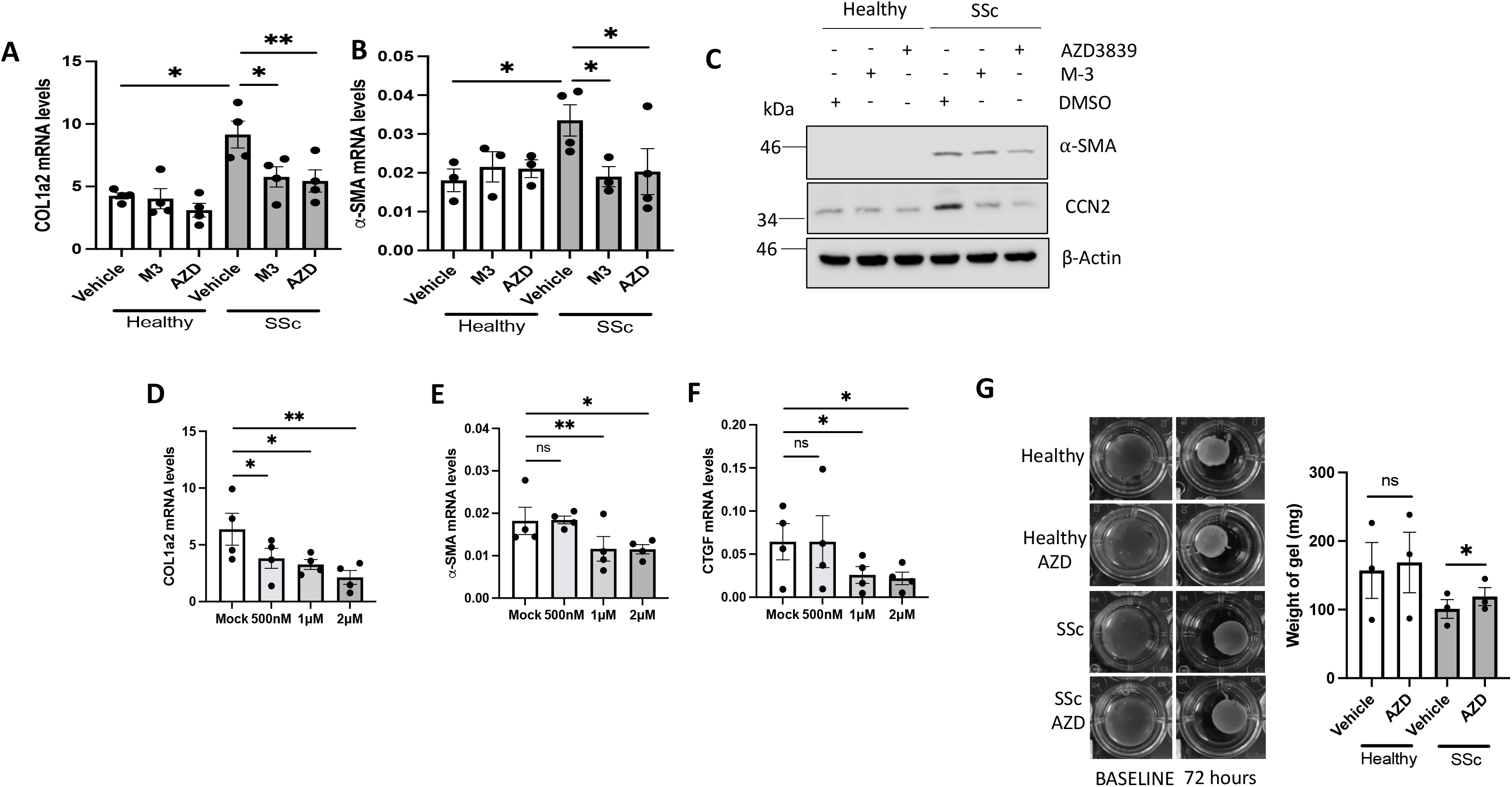
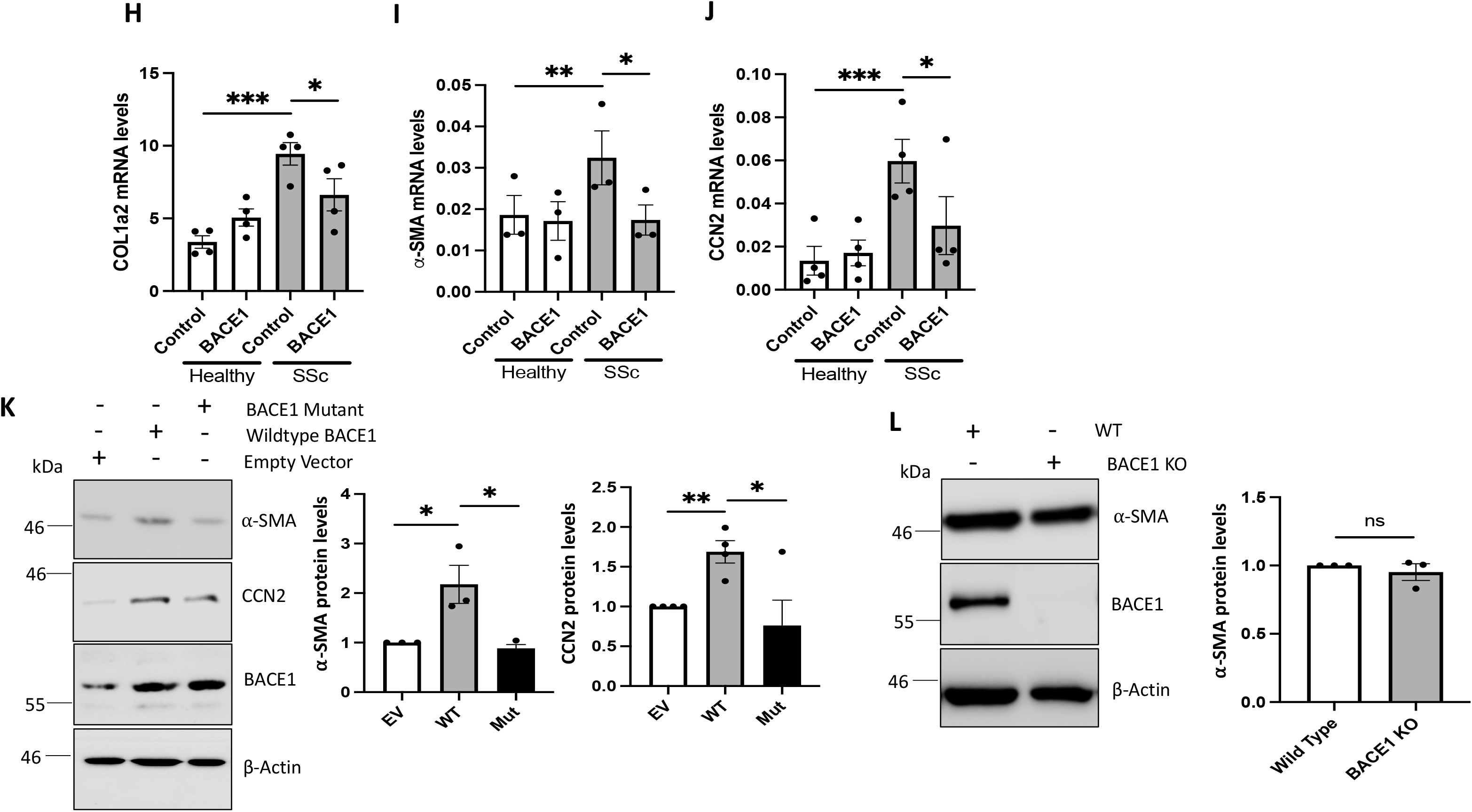
Modulation of BACE1 affects SSc fibroblast activation. RNA was extracted from healthy and SSc fibroblasts. In addition fibroblasts were treated with the BACE1 inhibitors M-3 (1µM) and AZD3839 (1µM) (A-C) or AZD3839 (500nM to 2uM) (D-F) or BACE1 siRNA (H-J) for 48 hours. Col1a2, α-SMA and CCN2 protein and transcript levels were assessed. (G) Healthy and SSc fibroblasts were suspended in collagen gels in the presence or absence of AZD3839 (1μM). Images were obtained for each condition and the weight of the gels were measured. (K) Healthy fibroblasts were transfected with mammalian vectors containing WT BACE1 or BACE1 mutant. (L) Fibroblasts were isolated from skin biopsies of WT and BACE1 KO mice (N=3 per group). Graphs represent the mean and standard error. *p<0.05, **p<0.01, ***p<0.001

Importantly, we observed similar results when we knocked down BACE1 expression using siRNA. BACE1 knockdown reduced Col1a2 (Figure2H), α-SMA (Figure2I) and CCN2 (Figure2J) transcript levels in SSc fibroblasts to similar levels seen in healthy fibroblasts. BACE1 knockdown in primary SSc fibroblasts resulted in a reduction in CCN2 protein expression (Supplementary Figure 2C)

In a complementary approach, we overexpressed BACE1 in healthy dermal fibroblasts to determine if elevated BACE1 protein expression is sufficient to drive fibroblast activation. Healthy dermal fibroblasts were transfected with mammalian expression vectors encoding wildtype (WT) BACE1 or an inactive mutant (D92A/D289A; Mut) (7) (Figure2K). While both BACE1 proteins expressed at similar levels (WT=3.6, Mut=3.7-fold increase in expression over endogenous levels in empty vector-transfected cells), only the wildtype BACE1 protein increased α-SMA and CCN2 expression at protein level in healthy fibroblasts. This confirms that increased levels of BACE1 can trigger fibroblast activation and that its proteolytic activity is essential for this function.

Next, we isolated skin fibroblasts from WT and BACE1 KO mice to explore the role of BACE1 in basal pro-fibrotic marker expression. We observed no differences in α-SMA protein levels between the wildtype and BACE1 KO dermal fibroblasts (Figure2M). This agrees with the data showing no effect of BACE1 inhibition or BACE1 siRNA-mediated knockdown on α-SMA in healthy fibroblasts transfected with the BACE1 siRNA. This suggests BACE1 requires fibrotic stimuli in order to regulate pro-fibrotic gene expression.

As BACE1 inhibition had no effect on basal pro-fibrotic marker expression in healthy fibroblasts, we tested if BACE1 is necessary for fibroblasts activation in response to defined pro-fibrotic stimuli. TGF-β, Wnt and Hh signalling are dysregulated in SSc fibroblasts and play an important role in SSc disease progression (2, 3, 4). The pro-fibrotic stimuli enhance the expression of several pro-fibrotic genes in fibroblasts, including α-SMA (3). Stimulation with either TGF-β, Wnt-3a or SAG (agonist of the Hh pathway) led to healthy fibroblasts activation (increased α-SMA levels) but had no effect on BACE1 levels (Figure3A). Next, we investigated the potential role of BACE1 as a downstream mediator of TGF-β, Wnt3a and SAG-induced fibroblast activation. To test this, we stimulated healthy fibroblasts with TGF-β in combination with each BACE1 inhibitor. Both BACE1 inhibitors blocked the TGF-β-mediated increase in α-SMA protein (Figure3B). Similar results were observed using BACE1 siRNA (Figure3C). Silencing of BACE1 attenuated TGF-β ability to increase α-SMA levels in dermal fibroblasts. This was also the case for Wnt3a-mediated fibroblast activation (Figures3D-E). Finally, SAG-dependent fibroblast activation was also sensitive to BACE1 inhibition and BACE1 siRNA (Figure3F-G). Together, these data suggest that BACE1 is an important downstream regulator of fibroblast activation when fibroblasts are challenged with pro-fibrotic stimuli.

### BACE1 activity is required for **β**-catenin stabilisation by Wnt3a

Next, we sought to determine the mechanism by which BACE1 regulates mediates the profibrotic effect of Wnt3a, TGF-β and Hh signalling. The well-established BACE1 substrate amyloid precursor protein (APP) has been recently shown to act as a Wnt decoy receptor, binding Wnt3a and Wnt5a through its cysteine-rich domain (18). We hypothesised that increased processing of APP by BACE1 in SSc as a consequence of higher expression levels could increase the availability of Wnt ligands for Frizzled-dependent activation of canonical Wnt/β-catenin signalling. In support, inhibition of BACE1 with AZD3839 or M3 reduced β-catenin protein levels in SSc fibroblasts, 30% and 15% respectively, but had no effect on protein levels in healthy control fibroblasts (Figure4A). These results were replicated in primary SSc dermal fibroblasts treated with M3 (Supplementary Figure 2B) In addition, we observed a dose-dependent decrease in β-catenin expression levels with AZD3839 in SSc fibroblasts (Supplementary-Figure 3A). BACE1 siRNA reduced β-catenin in immortalised and primary SSc fibroblasts (Figure 4B and Supplementary Figure 2C). We also investigated if BACE1 could also modulate β-catenin stability in response to exogenous Wnt3a. Healthy fibroblasts stimulated with Wnt-3a showed increased levels of β-catenin (Figure4C-D), which was reversed in the presence of the BACE1 inhibitors AZD3839 and M3. Overexpression of BACE1 in healthy dermal fibroblasts increased basal β-catenin expression levels (Figure4E) in an activity dependent manner, as the BACE1 inactive mutant was unable to increase β-catenin expression levels.

**Figure 3:**
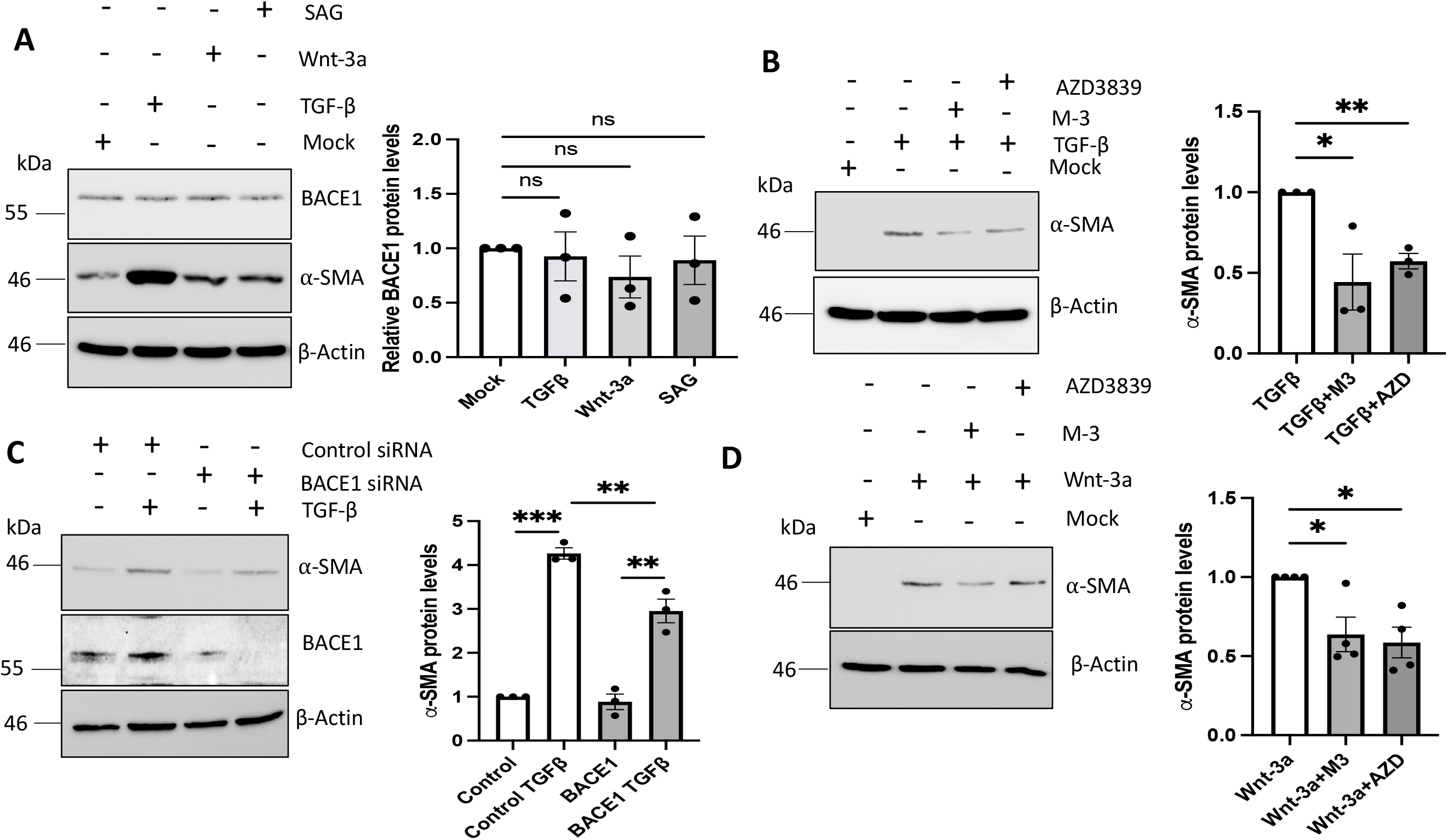
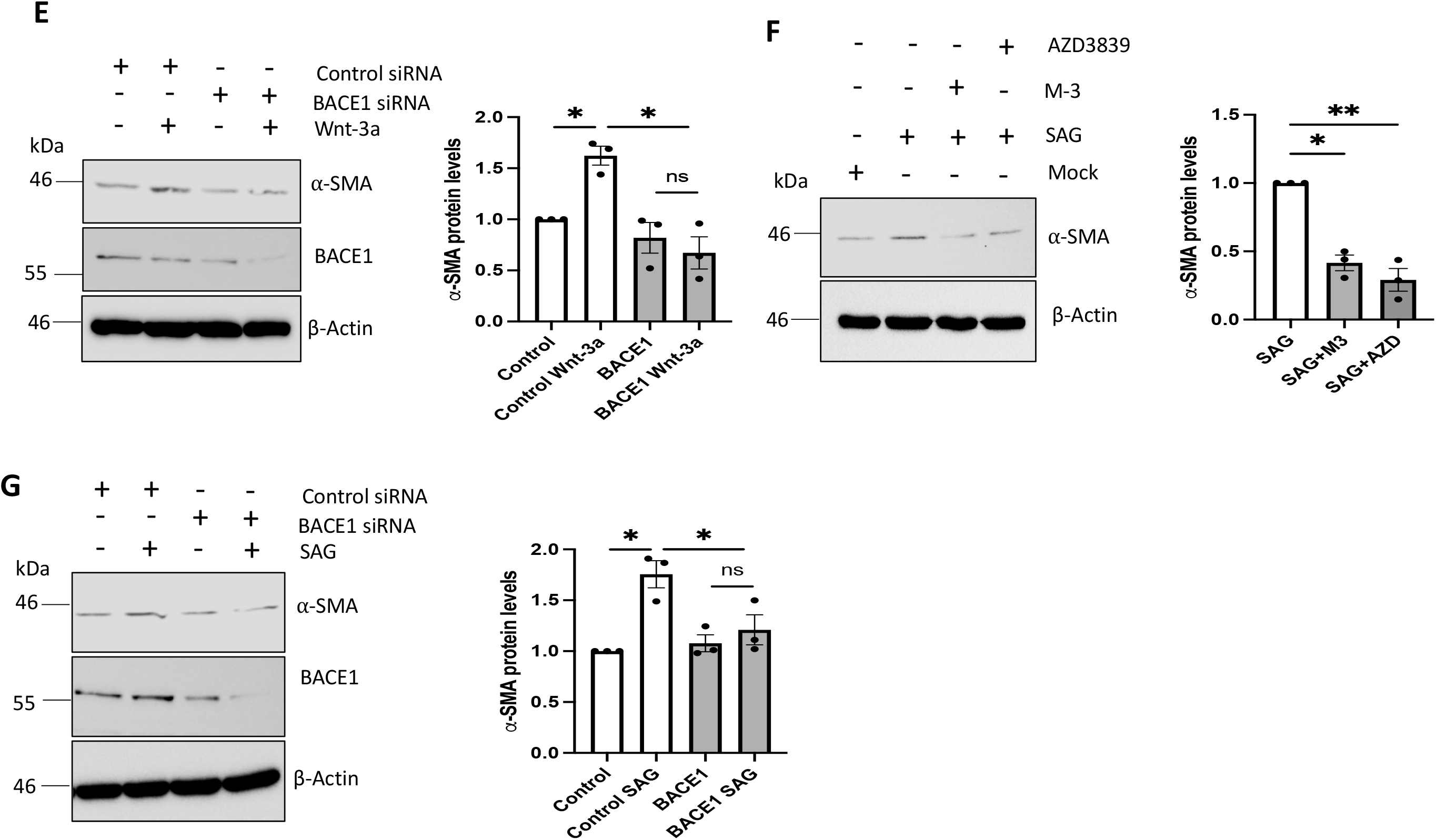
BACE1 is important for morphogen mediated fibroblast activation. Healthy dermal fibroblasts were grown in serum depleted media and stimulated with (A,B,C) TGF-β, (A,D,E) Wnt-3a or (A,F,G) SAG for 48 hours. BACE1 and α-SMA protein levels were assessed by western blot. β-actin was used as a loading control. In addition healthy dermal fibroblasts were treated with the BACE1 inhibitors M-3 or AZD3839 for 48 hours (B,D,F) or transfected with scramble or BACE1 specific siRNA for 48 hours (C,E,G). Graph represents densitometry analysis of the α-SMA western blots. *p<0.05, **p<0.01, ***p<0.001

**Figure 4:**
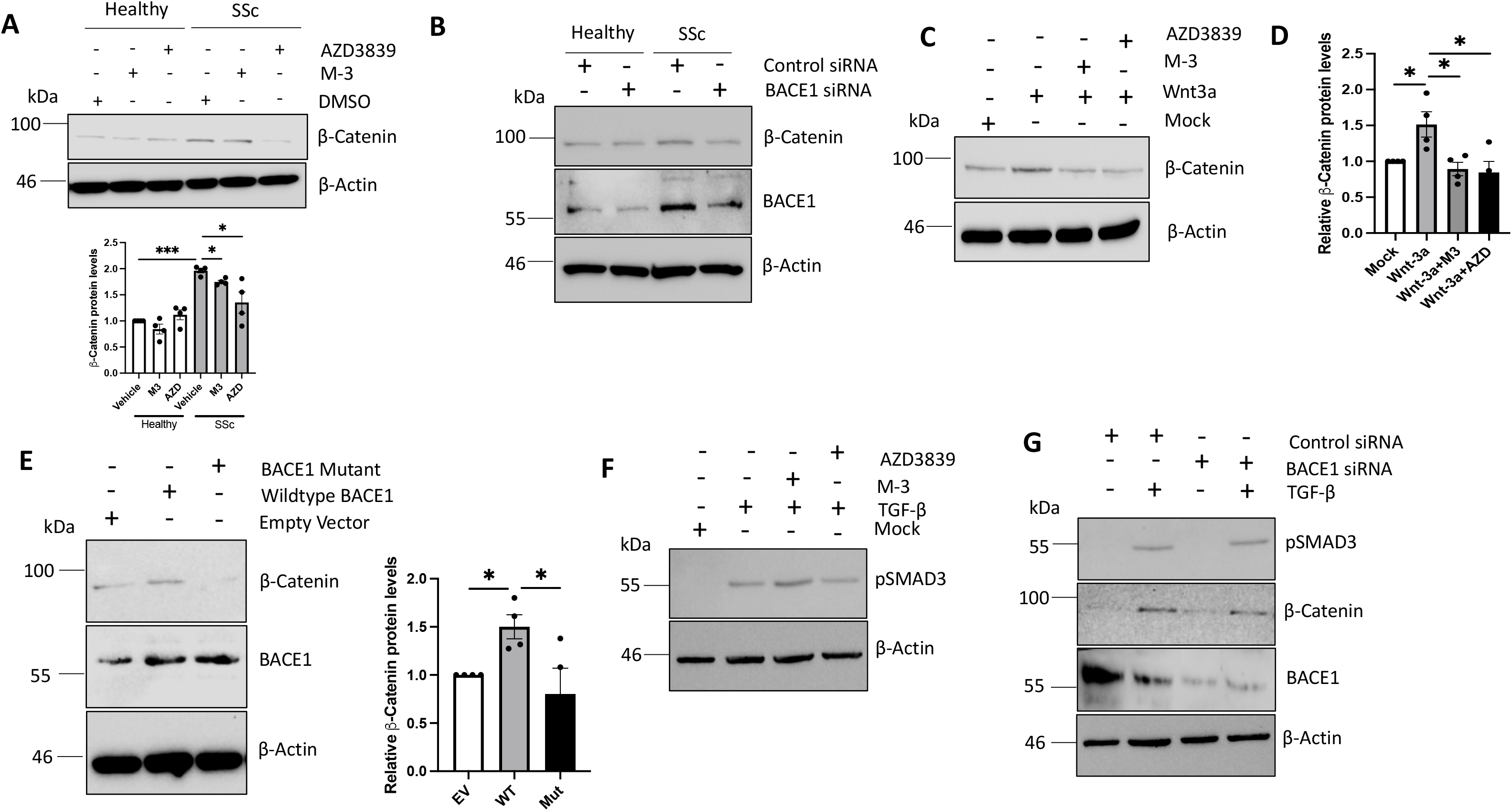
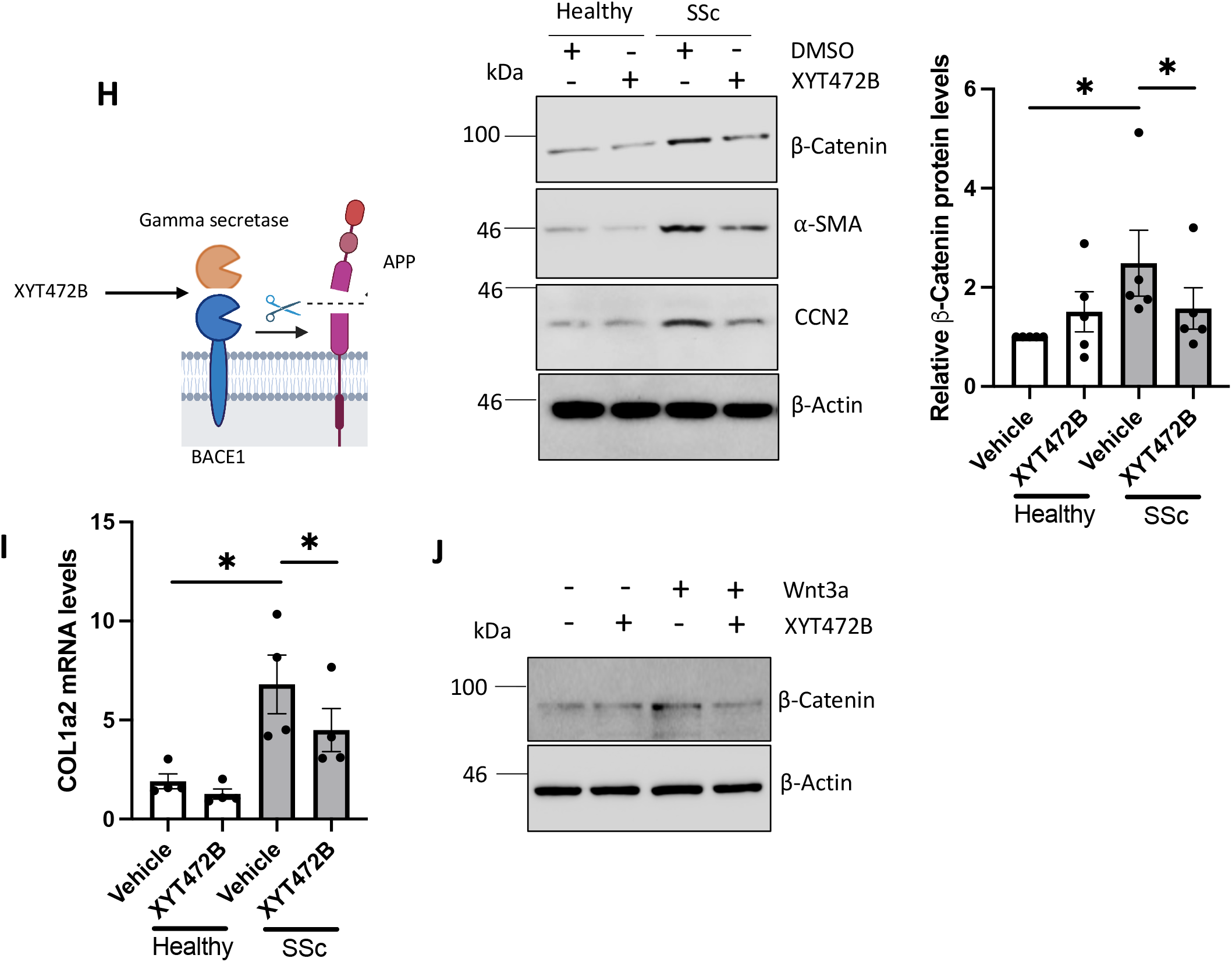
BACE1 regulates β-catenin expression in the presence and absence of Wnt ligands. Protein was extracted from healthy and SSc fibroblasts. In addition the fibroblasts were treated with the BACE1 inhibitor AZD3839 (AZD) and M3 (A), BACE1 siRNA (B). Healthy fibroblasts were stimulated with Wnt-3a (C, D, J) or TGF-β (F) for 48 hours in the presence and absence of BACE1 inhibitors. (E) Healthy fibroblasts were transfected with mammalian vectors containing WT BACE1 or BACE1 mutant. (G) Healthy fibroblasts were transfected with scramble or BACE1 specific siRNA and stimulated with TGF-β for 48 hours. (H-I) RNA and protein were extracted from healthy and SSc fibroblasts. In addition, healthy and SSc fibroblasts were treated with the BACE1 inhibitors XYT472B (10µM)) for 48 hours. *p<0.05, **p<0.01, ***p<0.001

Previous studies have shown that β-catenin is an important mediator of TGF-β-mediated, as inhibition of β-catenin with the small molecule inhibitor FH535 blocked TGF-β ability to induce α-SMA expression (3). In order to determine if the role of BACE1 in fibroblast activation by TGF-β is solely dependent on β-catenin or if it also regulates the TGF-β/SMAD3 axis, we studied SMAD3 phosphorylation. Interestingly, Inhibition of BACE1 (Figure4F) or BACE1 knockdown (Figure4G) had no effect on phosphorylation of SMAD3 in response to TGF-β stimulated healthy fibroblasts. Here we show TGF-β stimulation increases β-catenin levels in dermal fibroblasts and this was partially blocked with the BACE1 siRNA (Figure4G). This suggests BACE1 regulates the TGF-β signalling pathway downstream of SMAD3 through the modulation of β-catenin.

Having demonstrated that BACE1 activity is necessary for β-catenin increase in response to Wnt3a, we tested our hypothesis that BACE1 promotes Wnt signalling by proteolysis of full-length APP. To test this hypothesis, we used XYT472B an inhibitor that specifically prevents BACE1 from processing APP, but importantly does not affect the ability of BACE1 to cleave other substrates (19). Following treatement of healthy and SSc fibroblasts with the compound (Figure4H-J). We observed a reduction in β-catenin protein levels in SSc fibroblasts treated with XYT472B to levels similar to healthy control fibroblasts (Figure4H). As expected, the reduction in β-catenin levels resulted in reduced α-SMA, CCN2 protein levels (Figure 4I) and Col1a2 transcript levels (Figure4J) transcription in SSc fibroblasts treated with XYT472B to similar levels as the other inhibitors (Figure2). In addition, XYT472B blocks Wnt3a mediated induction of β-catenin expression. Healthy fibroblasts stimulated with Wnt-3a had increased β-catenin expression and this was attenuated with the addition of XYT472B (Figure4J). This data suggests BACE1 modulates canonical Wnt signalling through processing of APP, a Wnt decoy receptor.

### BACE1 regulates myofibroblast activation through the activation of Notch1 signalling

The Notch signalling pathway has been implicated in SSc dermal fibrosis (20) and there is evidence that the pathway is regulated by BACE1 (21). The Notch receptor resides at the cell surface and upon ligand binding (Jagged-1 (JAG1) or Delta like ligand-1 (DLL1)), it is cleaved to release a soluble the active Notch intracellular domain (NICD), which induces transcription of downstream target genes (*HES1/HEY1*). In SSc fibroblasts, Notch signalling is enhanced as assessed by increased NICD and *HES1* expression. Generation of NICD is orchestrated by consecutive action of an extracellular protease and the gamma secretase enzyme complex and inhibition of gamma secretase blocks SSc fibroblast activation (20). To explore the possibility that BACE1 contributes to Notch1 processing in SSc, we assessed Notch signalling in SSc fibroblasts treated with the BACE1 inhibitors. Notch1 expression was upregulated in SSc fibroblasts but the BACE1 inhibitors had no impact on Notch1 transcript levels (Figure5A). Nevertheless, Hes1 transcript levels (downstream transcriptional target of Notch) were significantly reduced in SSc fibroblasts treated with the BACE1 inhibitors in a dose dependent manner (Figure5B, Supplementary-Figure3B). In the same experimental setting, Hes1 levels of healthy fibroblasts were not affected by the inhibitors. The BACE1 inhibitors reduced the levels of NICD in SSc fibroblasts in a dose dependent manner (Figure5C-D, Supplementary-Figure3 A, C). This suggests BACE1 regulates the activation of Notch signalling in SSc fibroblasts through the activation of the receptor. The hedgehog transcription factor GLI2, has previously been shown to be a downstream target of the Notch pathway in SSc fibroblasts (4). The BACE1 inhibitors reduced the expression of the Hedgehog transcription factor GLI2 in SSc fibroblasts (Figure5C). Consistent with this data, BACE1 knock down by siRNA did not alter Notch1 (Figure5E) transcript levels, but did reduce Hes1 (Figure5F) and GLI2 (Figure5G) transcript levels, as well as NICD and GLI2 protein levels (Figure5H). In a complementary, gain of function set of experiments, overexpression of BACE1 resulted in increased accumulation of NICD (Figure5I), while overexpression of the secretase mutant was unable to induce NICD accumulation.

Interestingly, overexpression of BACE1 reduced full length Jagged1 protein levels in dermal fibroblasts (Figure5I). BACE1 has been shown to selectively regulate the cleavage of membrane bound JAG1 in Schwann cells (21). Regulation of JAG1 by extracellular domain shedding may play a role in the activation of Notch 1 in SSc fibroblasts. Our data suggests that BACE1 cleaves Jagged 1 from the cell surface resulting in a soluble JAG1 (sJAG1), which is enough to induce Notch signalling in a paracrine signalling cascade. The reduction in JAG1 levels was not observed in fibroblasts overexpressing the secretase dead mutant (Figure 5I). Further analysis of JAG1 levels in healthy and SSc fibroblasts treated with the BACE1 inhibitors revealed low levels of JAG1 in SSc fibroblasts and this was partially reversed with both AZD3839 and M3 (Figure5C). This suggests that BACE1 might cleave JAG1 from the surface of SSc fibroblasts which in turn could lead to increased soluble jagged 1 that is able to induce Notch signalling.

**Figure 5:**
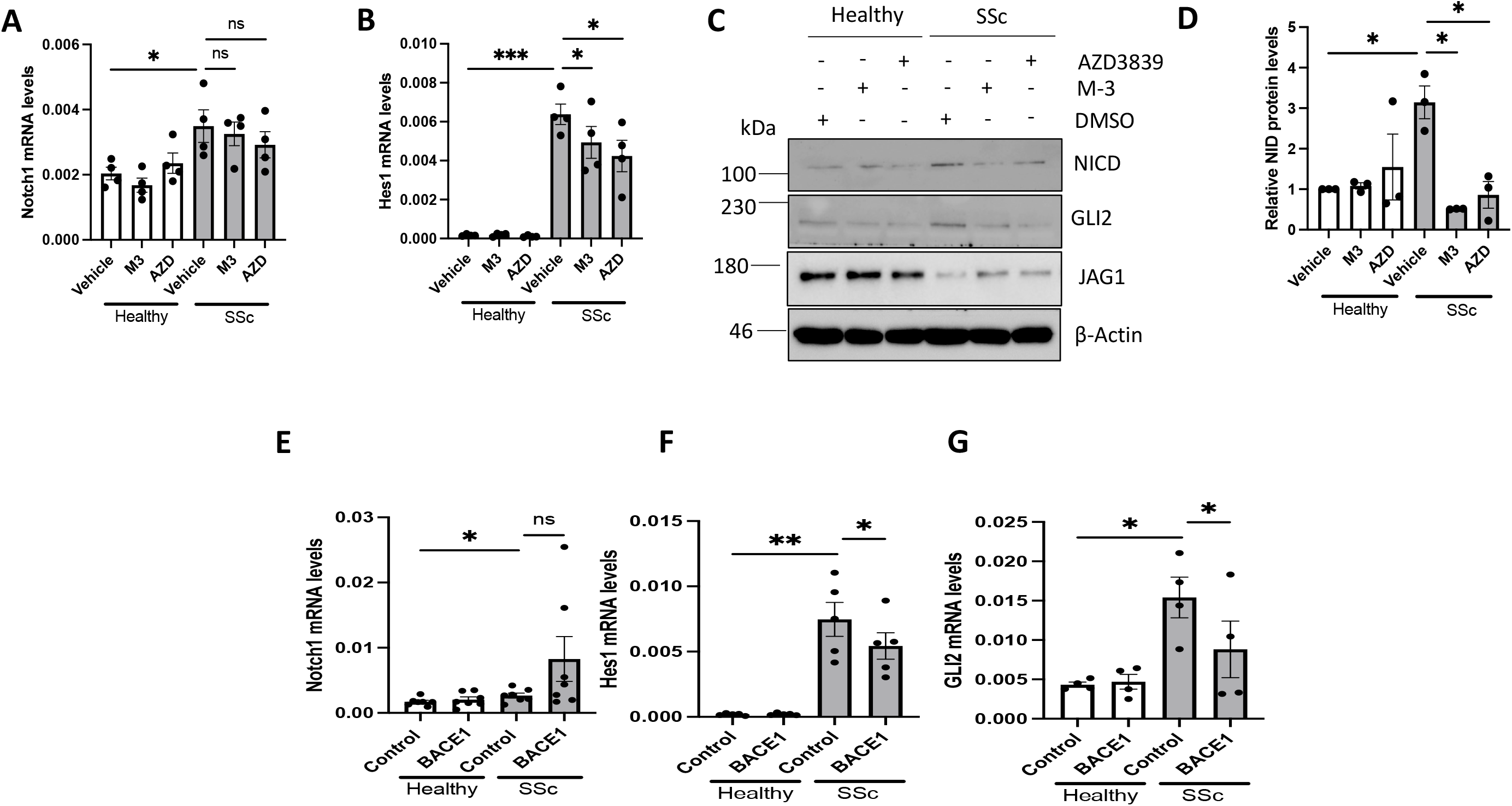
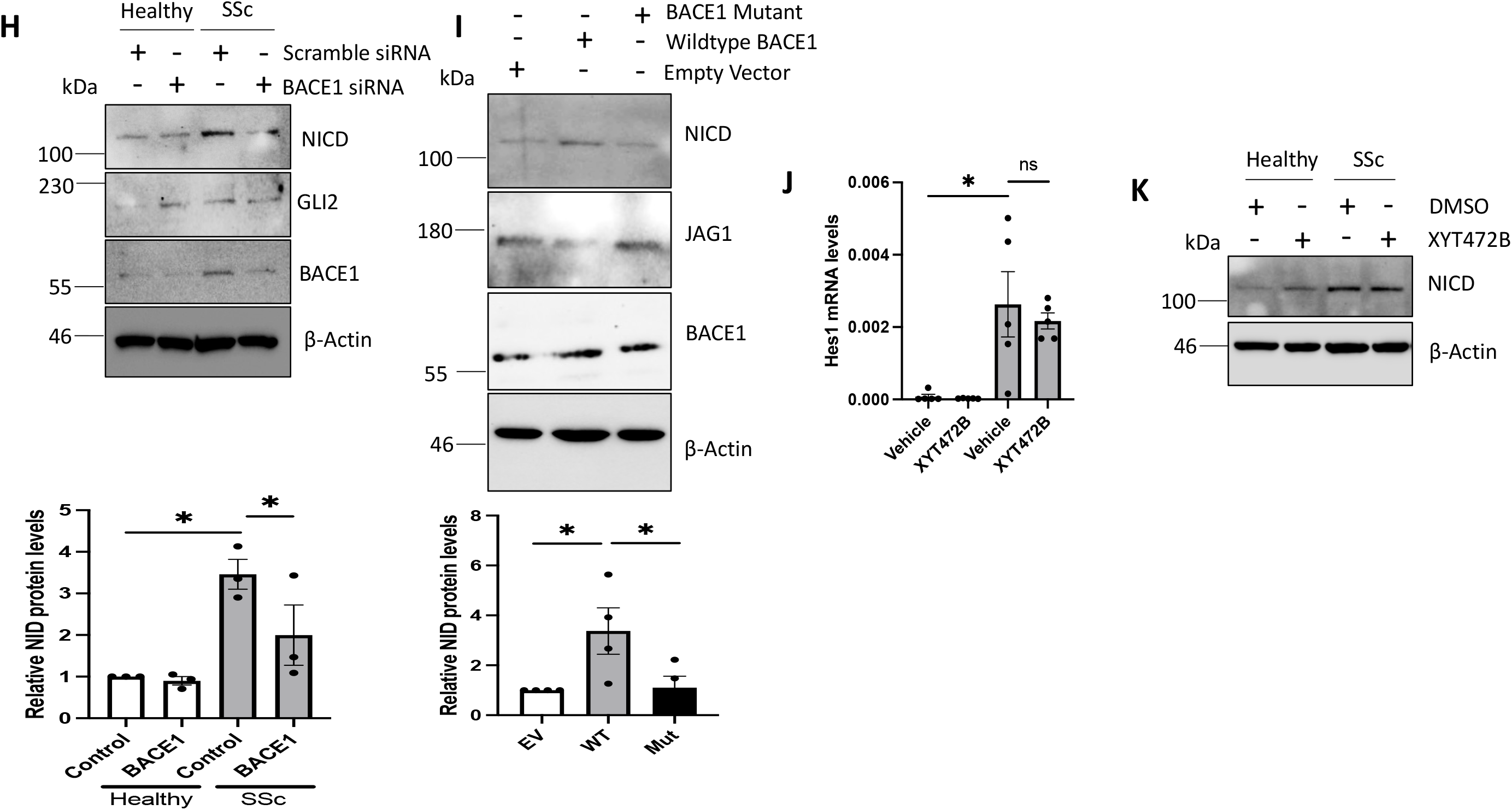
Inhibition of BACE1 leads to reduced Notch signalling in SSc fibroblasts. Healthy and SSc fibroblasts were treated with the BACE1 inhibitors M-3 and AZD3839 (A-D), BACE1 siRNA (E-H). Notch 1 and Hes1 transcript levels were assessed by qPCR and NICD, GLI2 and Jagged 1 protein levels were assessed by western blot. (I) Healthy fibroblasts were transfected with mammalian vectors containing WT BACE1 or BACE1 mutant. BACE1, NICD and Jagged 1 protein levels were assessed by western blot. β-actin was used as a loading control. RNA was extracted from healthy and SSc fibroblasts. In addition healthy and SSc fibroblasts were treated with the BACE1 inhibitors XYT472B (10µM)) for 48 hours. Hes1 (J) transcript levels were assessed by qPCR and NICD protein levels (K). *p<0.05, **p<0.01, ***p<0.001

Next, we assessed the effects of the APP-specific BACE1 inhibitor XYT472B on Notch signalling. XYT472B, was unable to inhibit or Hes1 transcription (Figure5J) or NICD expression (Figure 5K) in SSc fibroblasts. This suggests BACE1 ability to induce the Notch signalling cascade, is independent from its ability to target APP.

Further investigations found that inhibition or silencing BACE1 reduced the ability of TGF-β, Wnt and Hh signalling to modulate NICD in healthy fibroblasts (Supplementary Figure4A-B). We observed increased NICD protein levels in TGF-β stimulated fibroblasts, which was reversed when the fibroblasts were treated with M3, AZD3839 (Supplementary Figure4A) or BACE1 siRNA (Supplementary Figure4B). Cooperation between TGF-β Notch signalling pathways is well characterised with Notch signalling required for the regulation of a number of TGF-β responsive genes (22). Similar results were observed in Wnt3a (Supplementary Figure4C-D) and SAG (Supplementary Figure4E-F) stimulated fibroblasts. Together our data shows BACE1 regulates Notch signalling in SSc fibroblasts, which in turn contributes to fibroblast activation.

### BACE1 expression and pro-fibrotic function are attenuated by BDNF

We have identified that BACE1 is upregulated in SSc dermal fibroblasts. To date, the mechanism behind the elevated BACE1 expression levels is unresolved. A number of pathways known to regulate BACE1, in other diseases and tissues, are dysregulated in SSc fibroblasts. Therefore, we aimed to determine the pathway(s) involved in regulating BACE1 expression in SSc fibroblasts.

First, a number of potential mediators were excluded. Overexpression of epigenetic factors such as the lncRNA HOTAIR (20) in dermal fibroblasts did not affect BACE1 levels (Supplementary Figure 5A). Oxidative stress is dysregulated in SSc (23, 24) and has been shown to regulate BACE1 in neuronal cells (25). Induction of oxidative stress in dermal fibroblasts increased BACE1 transcription but not protein levels (Supplementary Figure 5B-C). BACE1 is known to interact with the lipid raft protein caveolin 1 (Cav-1) (26) and we have previously shown Cav-1 is downregulated in SSc fibroblasts (27, 28). Knockdown of Cav-1 in dermal fibroblasts reduced BACE1 levels (Supplementary Figure 5D). These data suggest other factors are at play in the overexpression of BACE1 in SSc fibroblasts.

Previous studies have shown the Brain Derived Neurotropic Factor (BDNF) plays an important role in regulating the activity of BACE1 (29). BDNF was shown to suppress BACE1 activity in mouse neuronal cells (29). To determine if BDNF regulates BACE1 expression and activity levels in dermal fibroblasts, we stimulated Healthy and SSc dermal fibroblasts with recombinant BDNF. Interestingly, BDNF stimulation reduced BACE1 expression and [-catenin levels in both healthy and SSc dermal fibroblasts (Figure6A). BDNF is known to signal through the Tropomyosin Receptor Kinase B (TRKB) in a range of cell types. Interestingly, there is evidence that shows that activation of TRKB attenuates liver fibrosis through inhibiting TGF-β signalling (30). Therefore, we wanted to determine if BDNF suppressed SSc fibroblast activation through its canonical receptor. Healthy dermal fibroblasts were stimulated with BDNF in combination with the TRKB inhibitor ANA-12. BDNF stimulation suppressed BACE1, β-catenin, α-SMA, CCN2 levels and increased JAG1 levels (Figure6B), and these effects were prevented by TRKB inhibition. This suggests BDNF alters BACE1 expression through its well characterised receptor TRKB.

**Figure 6:**
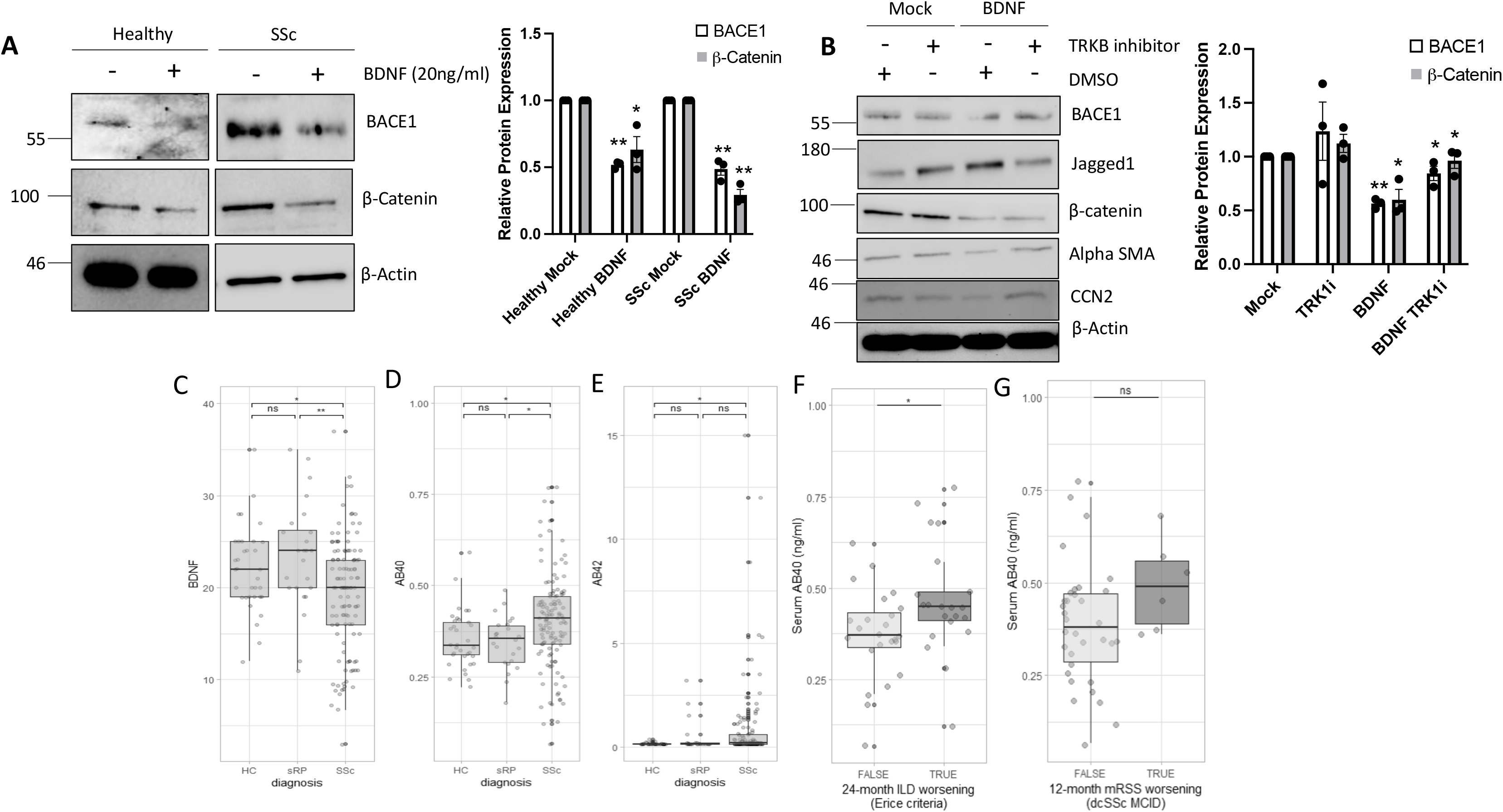
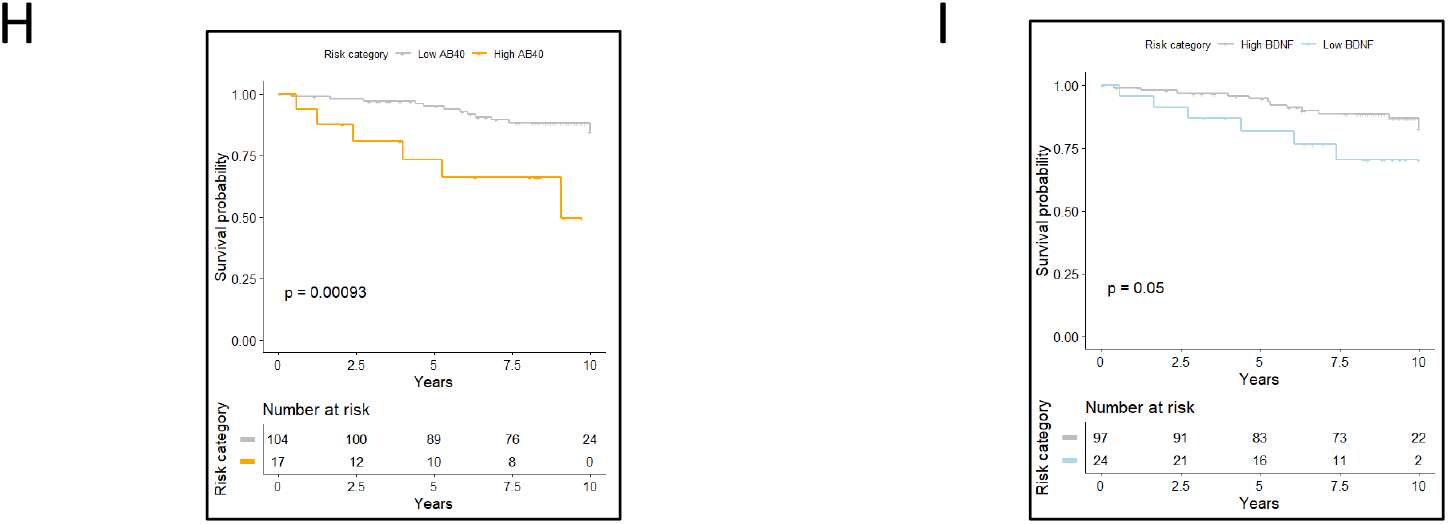
BACE1 expression levels are regulated by BDNF in SSc. Healthy and SSc fibroblasts were stimulated with BDNF (A) or in the presence of TRKB inhibitor (B). BACE1, Jagged1, α-SMA and β-catenin protein levels were assessed. Graphs represents densitometry analysis for both BACE1 and β-catenin western blots. Graph represents BDNF (C) AB40 (D) and AB42 (E) protein expression levels in Healthy (34), sRP (24) and SSc (121) patient sera. (F) Baseline serum AB40 levels in SSc-ILD patients according to the occurrence of Erice progression at 24-month follow-up. (G) Baseline serum AB40 levels in dcSSc patients according to occurrence of MCID mRSS worsening at 12-month follow-up. 10-year Survival probability charts of SSc patients categorized with high and low AB40 (H) and BDNF (I).

### Increased BACE1 activity markers and reduced BDNF levels in sera from SSc patients

As described above, BDNF is an important negative regulator of BACE1 in SSc patient fibroblasts. To investigate if the upregulated BACE1 activity in SSc could be linked to reduced BDNF, we assessed BDNF levels in sera samples from 121 SSc patients, 34 healthy controls (HC; aged 51.1±14.4 years, males 23.5%) and 24 subjects with secondary Raynaud’s phenomenon (sRP) (age 48.6±14.4 years, males 16.9%) via Luminex multiplex assays (Myriad RBM). Demographic and clinical characteristics of the involved SSc patients are reported in Supplementary Table1. BDNF levels were lower in SSc patients (19.20 ± 6.14 ng/ml) compared to both HCs (22.00 ± 4.61 ng/ml, adj. p-value=0.022), and sRP (23.40 ± 5.99 ng/ml, adj. p-value = 0.005) (Figure6C).

BACE1 cleaves APP to release soluble Aβ40 and Aβ42. Consistent with BDNF serum concentration data, serum Aβ40 levels were higher in SSc patients (0.40 ± 0.13 ng/ml) compared to both HCs (0.35 ± 0.08 ng/ml, adj. p-value=0.043) and sRP patients (0.35 ± 0.07 ng/ml, p-value= 0.043) (Figure6D). Similarly, Aβ42 was detectable in 56.4% of SSc patients compared to the 20.6% of HCs (adj. p-value=0.002; Figure6E). When the undetectable Aβ42 was approximated to the lower limit of detection, median Aβ42 values were higher in SSc patients [0.18 (IQR 0.14-0.59)] compared to HCs [0.14 (IQR 0.14-0.14), adj. p-value = 0.001] (Figure6E). Altogether, these data suggest that the low levels of circulating BDNF in SSc patient sera could underlie the increased BACE1 expression and activity in SSc.

### Serum A**β**40 and BDNF levels predict prognosis in SSc patients

In the subgroup of 45 patients with HR-CT proven ILD, 21 (46.7%) experienced a progression in the 24 months after serum collection according to Erice criteria (31). These progressor patients had higher baseline levels of Aβ40 (0.47 ± 0.15 ng/ml) compared to patients that were clinically stable over time (0.37 ± 0.12 ng/ml, p-value=0.028) (Figure6F). In the subgroup of 40 patients affected by diffuse cutaneous SSc variant, 6 (15.0%) experienced a clinically meaningful mRSS progression (32) in a 12-month period after serum collection. Also in this case, patients who experienced skin progression showed higher Aβ40 levels (0.49 ± 0.12 ng/ml) compared to the rest of the cohort (0.39 ± 0.16 ng/ml); however, statistical significance was not reached given the low number of progressors (p-value=0.099) (Figure6G).

Finally, 44 patients (36.4%) died during a 10-year follow-up and 19 (15.7%) of them as a direct consequence of cardio-pulmonary SSc involvement. For survival analysis, SSc patients were assigned to different risk categories according to their BDNF and Aβ40 levels compared to HCs. High-Aβ40 patients (14.0%) were defined are those with serum levels 2 standard deviations (SD) above the mean in the HC group and low-BDNF patients (19.8%) were those with serum levels 2 SD below the mean the HC group. The survival distributions of the defined risk categories were statistically different, with a poorer SSc-specific survival for high Aβ40 patients compared to low [χ2(2) = 11.0, p-value <0.001] (Figure6H) and for low BDNF patients compared to high [χ2(2) = 3.9, p-value = 0.05] (Figure6I).

## Discussion

This is the first study describing a role for BACE1 in SSc fibroblast activation. BACE1 levels are increased in SSc patient skin and isolated dermal fibroblasts. The elevated expression levels are important for SSc disease pathogenesis, as blocking BACE1 activity (with small molecule inhibitors) or expression levels (siRNA) led to a suppression of fibroblast activation in SSc. Moreover, the BACE1 products Aβ40 and Aβ42 were higher, and its negative regulator BDNF was lower, in SSc patient sera compared to HCs. From a clinical perspective, a status systemic fibroblast hyperactivation suggested by these markers identifies the patients with a poorer prognosis in terms of pulmonary and possibly skin fibrosis as well as overall mortality. The data presented in this study is preliminary and the longitudinal outcome data may be confounded by other baseline factors. Therefore further studies in validation cohorts will be needed to be performed to fully assess the potential of BDNF, Aβ40 and Aβ42 as predictive biomarkers in SSc.

The link between BACE1 and Alzheimer’s disease progression is interesting in the context of SSc. SSc patients with Alzheimer’s disease as a co-morbidity have a higher mortality rate (34). This could be a result of the higher levels of BACE1 (in patient skin) and β-amyloid (in SSc patient sera) which results in exacerbated Alzheimer’s disease progression.

To date most studies describing BACE1 functions have been performed in neuronal cell types in the context of Alzheimer’s disease. Therefore, this study uncovers novel functions for BACE1 in another disease setting and highlights the need to study BACE1 in other cell types and disease settings.

An intriguing element of the study is the uncovering of a role for BACE1 downstream of pro-fibrotic signalling (TGF-β, Wnt-3a and Hh). Inhibition of BACE1 via siRNA or small molecule inhibitors blocked TGF-β, Wnt3a and Smo-mediated fibroblast activation (Figure3). Interestingly APP is a conserved Wnt receptor (18). APP binds to Wnt3a and Wnt5a resulting in the recycling and stabilisation of APP. Thus the ability of BACE1 to process APP may enable more Wnt ligands to induce canonical Wnt signalling and fibroblast activation (Supplementary Figure 6). Cleavage of APP may result in its degradation which in turn would release the bound wnt ligands. In addition, previous studies have shown an interplay between BACE1 and GSK3β which is a member of the β-catenin destruction complex (34). We observed reduced β-catenin levels in Wnt3a stimulated fibroblasts when BACE1 was inhibited (Figure4C-D) supporting this hypothesis. Furthermore, a BACE1 inhibitor (XYT472B) (19) which prevents the ability of BACE1 to process APP but does not affect any other BACE1 substrate blocked pro-fibrotic gene expression in SSc dermal fibroblasts (Figure4H) and attenuated β-catenin accumulation in response to Wnt3a in healthy fibroblasts (Figure4J).

The ability of BACE1 to reduce fibroblast activation by the other signalling pathways (TGF-β and Hh), could be linked to its ability to modulate Wnt3a/β-catenin. We have previously shown co-operation between the Wnt3a and TGF-β signalling pathways in dermal fibroblasts as β-catenin inhibitors blocked the pro-fibrotic ability of TGF-β (3). Here we show that BACE1 does not regulate the activity of the TGF-β transcription factor SMAD3 (Figure4F-G). The ability of BACE1 to regulate Hh mediated fibroblast activation (Figure3F-G) could also be due to its ability to regulate β-catenin, since we previously showed thatWnt-3a/β-catenin induce expression of the Hh transcription factor GLI2 (3).

In addition, BACE1 promotes Notch signalling in response to TGF-β, Wnt3a and SAG (Supplementary Figure3). This suggests an integrated pathway involving BACE1, Notch and β-catenin involved in SSc fibroblast activation. β-catenin has previously been shown to induce the expression of the Notch ligand JAG1 (35). Therefore, β-catenin could drive Notch signalling by increasing ligand expression in SSc fibroblasts. One caveat to this is that the APP-specific BACE1 inhibitor XYT472B does not affect Hes1 mRNA levels in SSc fibroblasts, suggesting a different substrate, such as JAG1. BACE1 has been reported to cleave the Notch ligand JAG1 from the cell surface in neurons (21) and recombinant sJAG1 can trigger Notch signalling and fibroblast activation (36). In this study, we show that BACE1 reduces the level of full-length JAG1 in dermal fibroblasts (Figure 5).

Systemic sclerosis fibroblasts have previously been shown to share several similarities with cancer associated fibroblasts (37). Interestingly cancer associated fibroblasts secrete β-amyloid and this drives the formation of neutrophil extracellular traps (NETs) in the associated tissue. NETs are decondensed chromatin that the neutrophils extrude into the extracellular space and this has a microbicidal effect. BACE1 inhibition reduces β-amyloid secretion and in turn reduces NETs from neutrophils in the associated tissue (38). This has implications for SSc, as NETosis in patient plasma in the early stages of the disease correlates with vascular complications (39). This further highlights the potential role of BACE1 in multiple aspects of SSc disease progression.

We have shown BDNF reduces BACE1 expression levels in both healthy and SSc dermal fibroblasts in a TRKB-specific manner (Figure6). The role of BDNF in fibrosis is a conflicting story. Our data suggests that BDNF is inversely correlated with SSc progression and mechanistic studies in dermal fibroblasts point towards an anti-fibrotic function. However, BDNF was previously shown to have a pro-fibrotic role in idiopathic pulmonary fibrosis (IPF), suggesting a disease-specific role (40). Analysis of a published proteomic antibody microarray from healthy vs SSc sera revealed a similar downward trend in BDNF sera levels between healthy control and SSc patients with ILD and PAH (41). This strengthens our hypothesis that BDNF is anti-fibrotic in the context of SSc.

The translational potential of this study is underscored by number of small molecule inhibitors targeting BACE1, including AZD3839, that have progressed through clinical trials for the treatment of Alzheimer’s disease, indicating safety and good tolerance by the patients. This opens up the possibility that these compounds could be re-purposed for the treatment of SSc and provide a novel therapy which is desperately needed.

## Data Availability statement

Data are available in the article itself and its supplementary materials

## Acknowledgements

**C.W.W** is supported by Susan Cheney Scleroderma fellowship. **R.W** is supported by an MRC DiMeN DTP studentship. **F.D.G** is supported by the National Institute for Health Research (NIHR) Leeds Biomedical Research Centre (BRC). The views expressed are those of the author and not necessarily those of the NIHR or the Department of Health and Social Care. **P.J.M** is supported by the British Heart Foundation (FS/4yPhD/F/20/34130 and FS/18/38/33659) and the Biotechnology & Biological Sciences Research Council (BB/V014358/1). AZD3839 was kindly provided by AstraZeneca through their Open Innovation Program.

## Contributions

Designing Research Study: C.W.W, E.D.L., N.A.R.D.G, F.D.G, P.J.M

Conducting Experiments: C.W.W, E.C, R.L.R, G.M, K.A.W, B.C.R, C.A, R.W, J.B

Acquiring Data: C.W.W, E.D.L., E.C, R.L.R, K.A.W, B.C.R, C.A, R.W, J.B

Analysing Data: C.W.W, E.D.L, R.L.R, C.A, L.D.L, N.A.R.D.G, C.S.M, F.D.G, P.J.M

Writing Manuscript: C.W.W, E.D.L, F.D.G, P.J.M

All authors reviewed and approved the final version of the manuscript

## Ethics Approval and consent to participate

The study was approved by NRES committee North East-Newcastle & North Tyneside: REC Ref:15/NE/0211 to FDG. All participants provided written informed consent to participate in this study. Informed consent procedure was approved by NRES-011NE to FDG by the University of Leeds. PPL number for the mouse work is: PP2103311 issued by the University of Leeds

## Patient and Public Involvement

Patients or the public were involved in the design, or conduct, or reporting, or dissemination plans of our research

---

**Supplemetary Figure 1:**
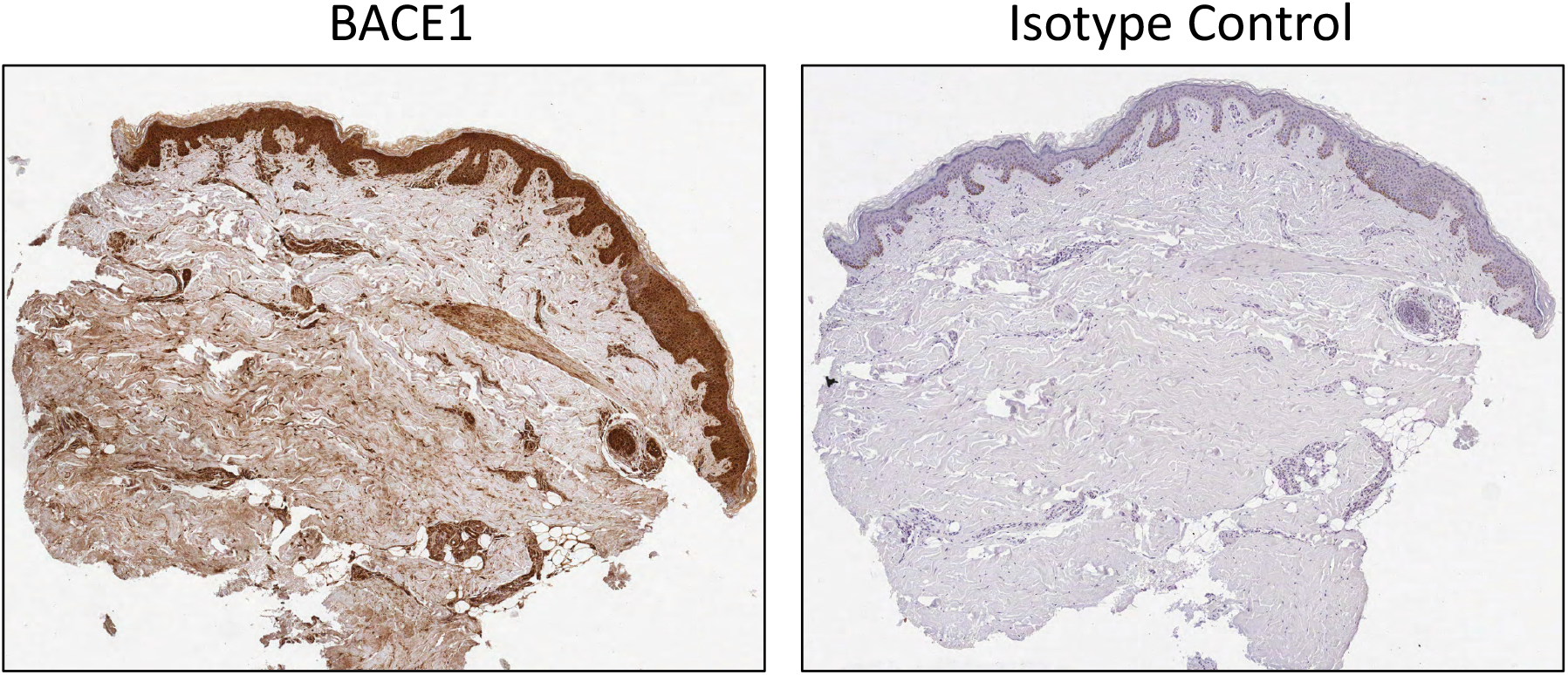
BACE1 levels are increased in SSc patient skin. SSc patient skin biopsies were stained with an antibody specific to BACE1 alongside a isotype control antibody. Skin sections were counterstained with Haematoxylin. Scale Bars represent 50μM

**Supplementary Figure 2:**
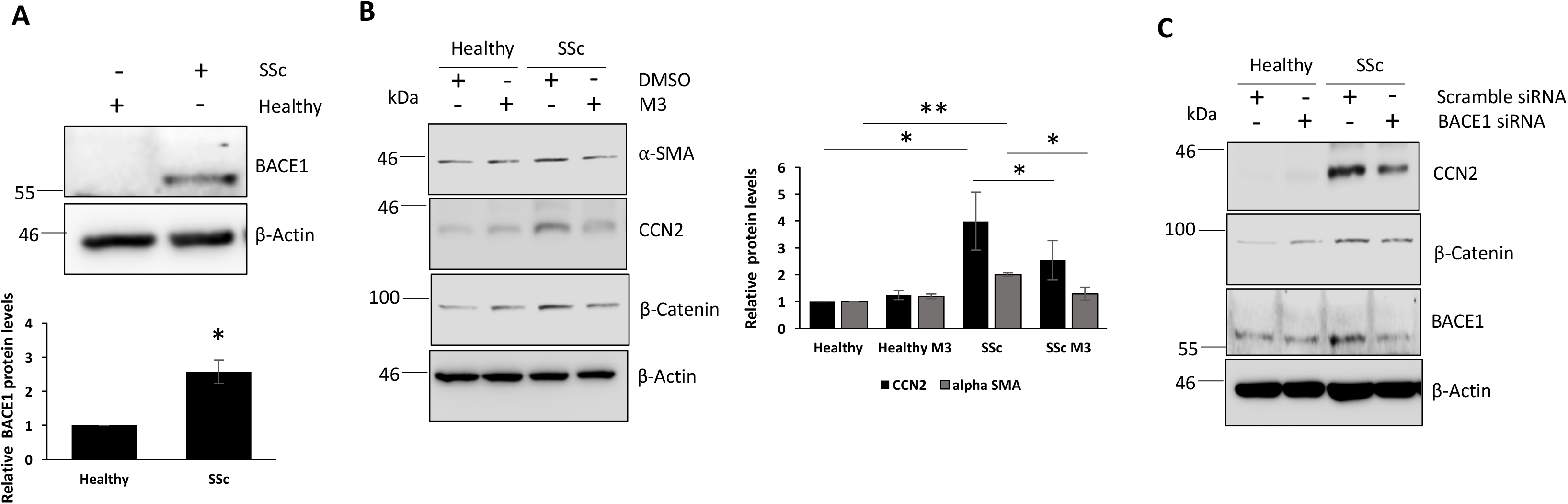
Modulation of BACE1 affects SSc fibroblast activation. (A)Protein was extracted from primary healthy and SSc fibroblasts. BACE1 protein levels were assessed. Protein was extracted from primary healthy and SSc fibroblasts. In addition fibroblasts were treated with the BACE1 inhibitors M-3 (1µM) (B) and BACE1 siRNA (C) for 48 hours. α-SMA, CCN2 and β-Catenin protein levels were assessed.

**Supplementary Figure 3:**
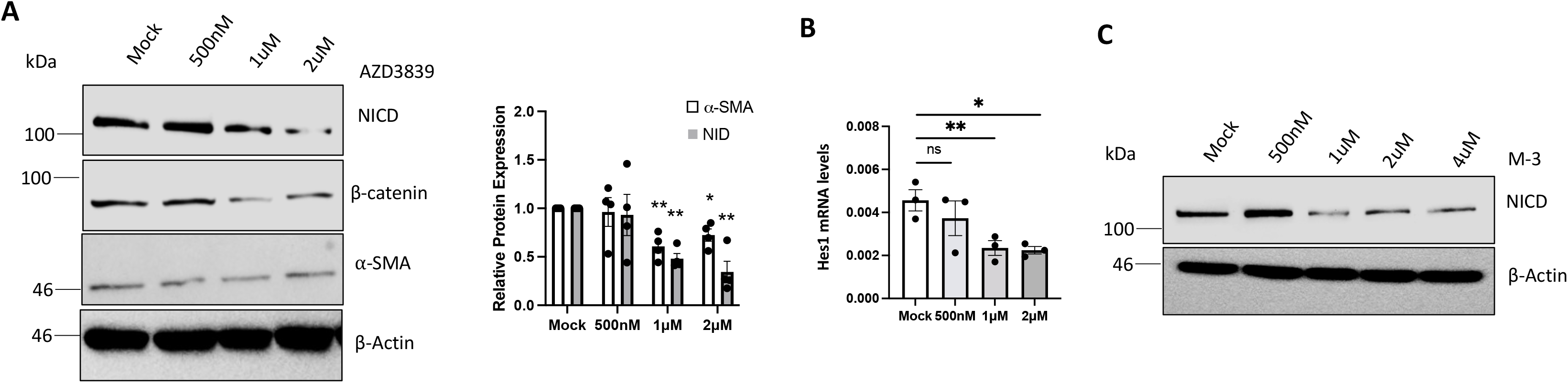
The BACE1 inhibitors block SSc fibroblast activation in a dose dependent manner. RNA and protein were extracted from SSc fibroblasts treated with the BACE1 inhibitors AZD (Range 500nM to 2uM) for 48 hours. (A) NICD, β-catenin and α-SMA protein levels were assessed by western blot. β-actin was used as a loading control. Graph represents densitometry analysis of western blots. (B) Hes1 transcript levels were assessed by qPCR. (C) Protein was extracted from SSc fibroblasts treated with the BACE1 inhibitors M3 (Range 500nM to 4uM) for 48 hours. NICD protein levels were assessed by western blot. β-actin was used as a loading control *p<0.05, **p<0.01, ***p<0.001

**Supplementary Figure 4:**
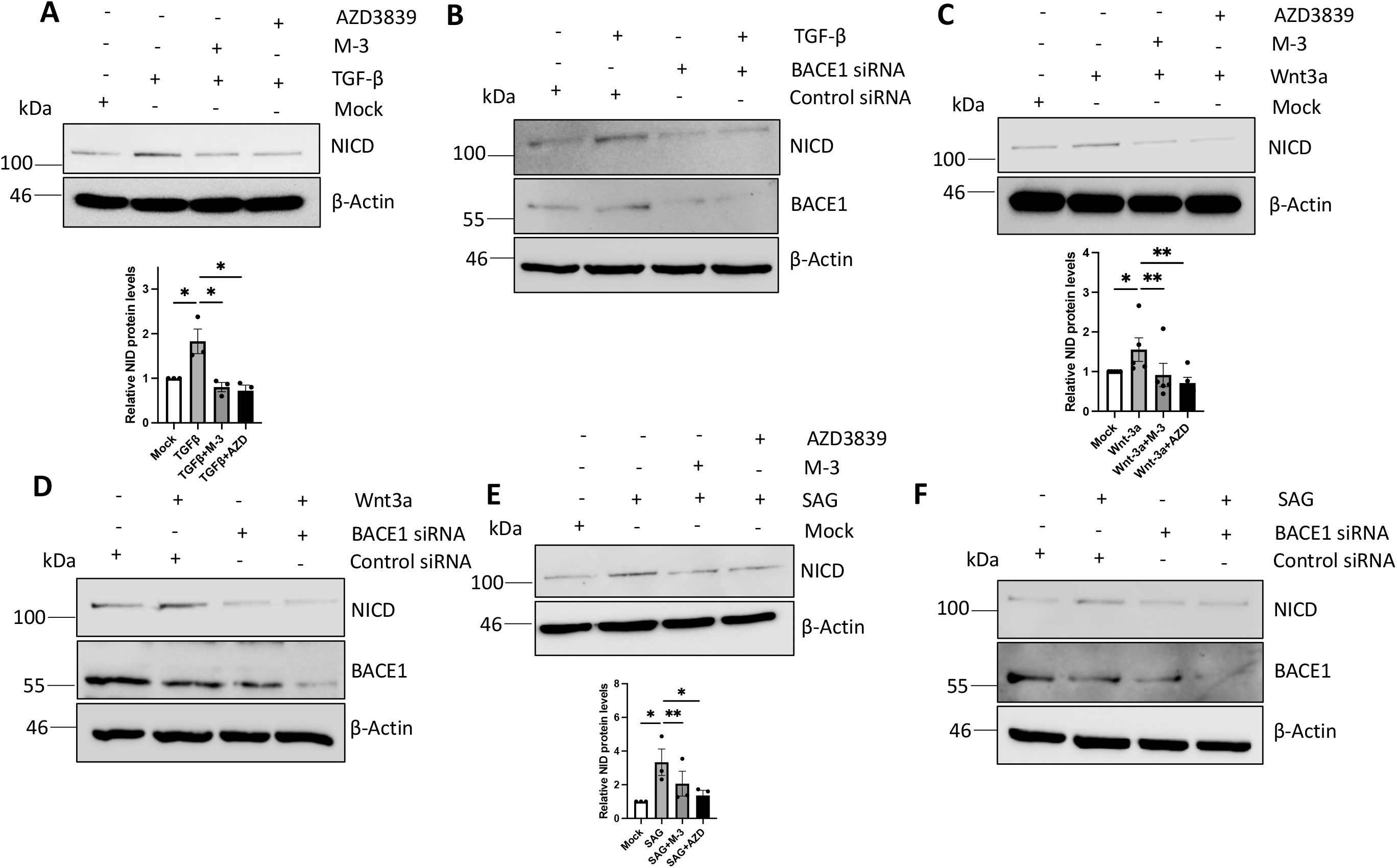
BACE1 regulates morphogen mediated Notch signalling. Healthy fibroblasts were grown in serum depleted media and stimulated with TGF-β (A), Wnt-3a (C) or SAG (E) for 48 hours in the presence or absence of BACE1 inhibitors, M-3 or AZD3839. NICD protein levels were analysed by western blot. β-actin was used as a loading control. Healthy dermal fibroblasts were transfected with scramble or BACE1 specific siRNA and stimulated with TGF-β (B), Wnt-3a (D) or SAG (F) for 48 hours. NICD protein levels were analysed by western blot. β-actin was used as a loading control. Graph represents densitometry analysis of NICD western blots. *p<0.05, **p<0.01, ***p<0.001

**Supplementary Figure 5:**
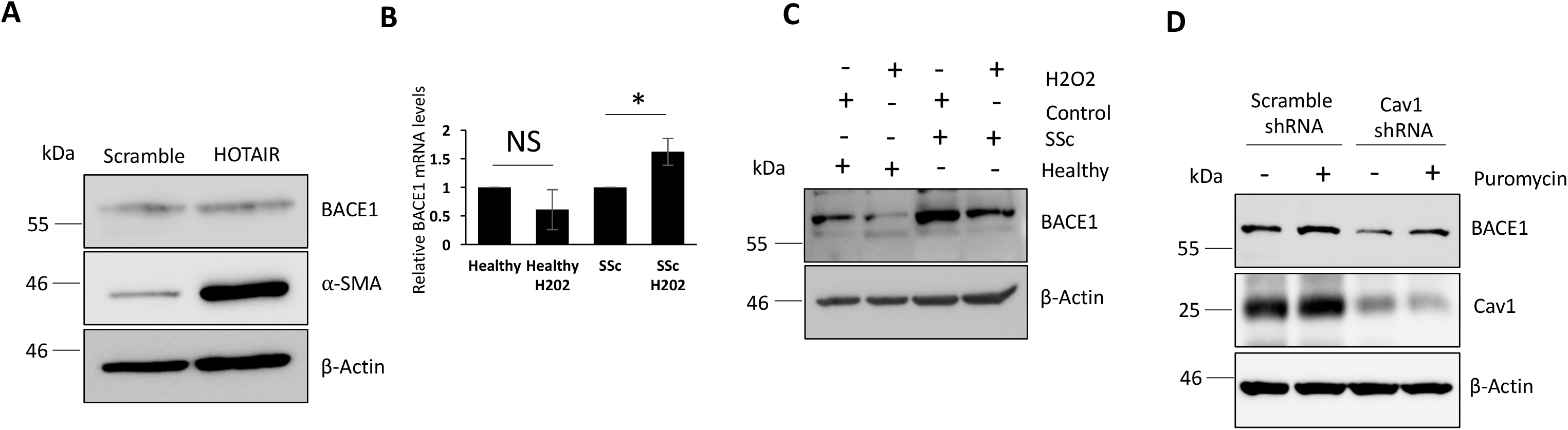
Regulation of BACE1 expression in dermal fibroblasts. Protein was extracted from fibroblasts expressing the sequence for HOTAIR or a scramble control. (A) BACE1 and α-SMA protein levels were assessed by western blot. RNA and protein were extracted from healthy and SSc dermal fibroblasts. In addition, the fibroblasts were treated with H_2_0_2_ for 24 hours. (B) BACE1 transcript levels were assessed by qPCR. Graphs represent the mean and standard error for three 3 independent experiments. (C) BACE1 protein levels were assessed by western blot. Protein was extracted from healthy dermal fibroblasts transduced with lentiviral vectors containing Cav-1 or scramble control shRNA. (D) BACE1 and Cav-1 protein levels were assessed by western blot.

**Supplementary Figure 6:**
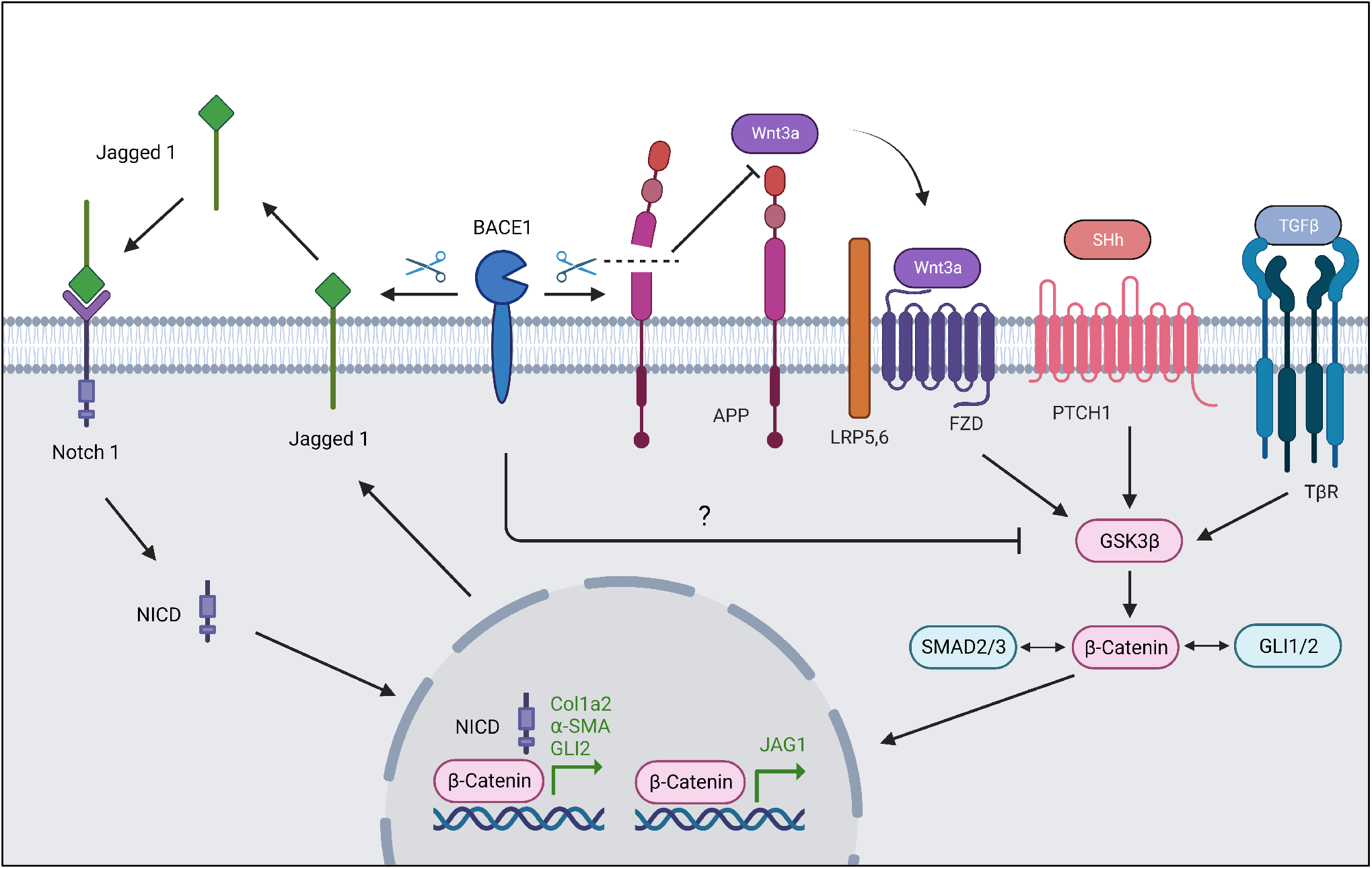
BACE1 regulates SSc fibroblast activation through a. β**-catenin/Notch axis.** BACE1 can regulate β-catenin expression through the processing of APP. BACE1 can activate Notch signalling through regulating the expression of the Notch ligand Jag1.

**Supplementary Table 1:** Demographic and disease characteristics of SSc patients evaluated for serum AB40, AB42 and BDN.

|  | SSc patients<br>N = 121 |
| --- | --- |
| Age, years, mean±SD | 57.4±12.5 |
| Male gender, n (%) | 21 (17.4) |
| Le Roy diffuse cutaneous variant, n (%) | 40 (33.1) |
| Disease duration, age, median (IQR) | 9.0 (4.0-14.0) |
| ACA, n (%) | 56 (46.3) |
| Anti-Scl70 antibody, n (%) | 27 (22.3) |
| Active-late capillaroscopy pattern, n (%) | 48 (64.9) |
| Late capillaroscopy pattern, n (%) | 26 (34.7) |
| mRSS, median (IQR) | 2.0 (1.0-5.0) |
| ILD on HRCT, n (%) | 45 (37.2) |
| PH on RHC, n (%) | 10 (8.3) |
| FVC, % of predicted, mean±SD | 100.5±23.6 |
| TLC, % of predicted, mean±SD | 91.8±17.8 |
| DLco, % of predicted, mean±SD | 62.8±16.1 |
| Digital ulcers, n (%) | 65 (53.7%) |
| Skin calcinosis, n (%) | 51 (42.1%) |
| Skin telangiectasias, n (%) | 94 (77.7%) |
| Arthritis, n (%) | 12 (9.9%) |
| Immunosuppressive treatment, n (%) | 42 (34.7%) |
| Vasoactive treatment, n (%) | 90 (74.4%) |
| AB40, ng/ml, mean±SD | 0.40±0.13 |
| AB42, ng/ml, median (IQR) | 0.18 (0.14-0.59) |
| BDNF, ng/ml, mean±SD | 19.20±6.14 |
| Abbreviations: AB amyloid β, ACA anticentromere antibody, BDNF Brain-derived neurotrophic factor, DLco lung diffusion of carbon monoxide, FVC forced vital capacity, HC healthy control, HRCT high-resolution computed tomography, IQR interquartile range, ILD interstitial lung disease, mRSS modified Rodnan skin score, RHC right heart catheterization, SD standard deviation, sRP secondary Raynaud's phenomenon, SSc systemic sclerosis, TLC total lung capacity. |  |

## References

1. Beyer C, Schramm A, Akhmetshina A, Dees C, Kireva T, Gelse K et al. β-catenin is a central mediator of pro-fibrotic Wnt signalling in systemic sclerosis. Ann Rheum Dis 2012;71(5):761–7

2. Gillespie J, Ross RL, Corinaldesi C, Esteves F, Derrett-Smith E, McDermott MF, et al. Transforming growth factor β activation primes canonical Wnt signalling through down-regulation of axin-2. Arthritis Rheumatol 2018;70:932–942

3. Wasson CW, Caballero-Ruiz B, Gillespie J, Derrett-Smith E, Mankouri J, Denton CP et al. Induction of pro-fibrotic CLIC4 in dermal fibroblasts by TGF-β/Wnt3a is mediated by GLI2 upregulation. Cells 2022;11(3):530

4. Wasson CW, Ross RL, Wells R, Corinaldesi C, Georgiou I Ch, Riobo-Del Galdo N et al. Long non-coding RNA HOTAIR induces GLI2 expression through Notch signalling in systemic sclerosis dermal fibroblasts. Arthritis Res Ther 2020;22:286

5. Hampel H, Vassar R, De Strooper B, Hardy J, Willem M, Singh N et al. The β-secretase BACE1 in Alzheimers’s disease. Biol Psychiatry 2021;89(8):745–756.

6. Taylor HA, Simmons KJ, Clavane EM, Trevelyan CJ, Brown JM, Przemlska L et al. PTPRD and DCC are novel BACE1 substrates differentially expressed in Alzheimer’s disease: A data mining and bioinformatics study. Int J Mol Sci 2022;23(9):4568

7. Meakin PJ, Mezzapesa A, Benabou E, Haas ME, Bonardo B, Grino M et al. The beta secretase BACE1 regulates the expression of insulin receptor in the liver. Nat Commun 2018;9(1): 1306

8. Del Galdo F, Hartley C, Allanore Y. Randomised controlled trials in systemic sclerosis: patient selection and endpoints for next generation trials. The Lancet Rheumatology 2020;2: e173–e184

9. Meakin PJ, Coull BM, Tuharska Z, McCaffery C, Akoumianakis I, Antoniades C et al. Elevated circulating amyloid concentrations in obesity and diabetes promotes vascular dysfunction. J Clin Invest 2020;130:4104–4117

10. Botteri G, Salvado L, Guma A, Hamilton DL, Meakin PJ, Montagut G et al. The BACE1 product sAPPβ induces ER stress and inflammation and impairs insulin signalling. Metabolism 2018;85:59–75

11. Heindryckx F, Binet F, Ponticos M, Rombouts K, Lau J, Kreuger J et al. Enoplasmic reticulum stress enhances fibrosis through IRE1α-mediated degradation of miR-150 and XBP-1 splicing. EMBO Mol Med 2016;8(7):729–44

12. Hendrik-Kuhn P, Marjaux E, Imhof A, De Strooper B, Haass C, Lichtenthaler SF. Regulated intramembrane proteolysis of the interleukin-1 receptor II by alpha-, beta- and gamma secretase. J Biol Chem 2007;282(16):11982–95

13. Iannazzo F, Pellicano C, Colalillo A, Ramaccini C, Romaniello A, Gigante A et al. Interleukin-33 and soluble suppression of tumorigenicity 2 in scleroderma cardiac involvement. Clin Exp Med 2022.

14. Chen C-H, Zhou W, Liu S, Deng Y, Cai F, Tone M et al. Increased NF-κB signalling up-regulates BACE1 expression and its therapeutic potential in Alzheimer’s disease. Int J Neuropsychopharmacol 2012;15(1):77–90

15. Wen W, Li P, Liu P, Xu S, Wang F, Huang JH. Post-translational modifications of BACE1 in Alzheimer’s disease. Curr Neuropharmacol 2022;20(1):211–222

16. Zhao Y, Zhou H, Zhao Y, Liang Z, Gong X, Yo J et al. BACE1 SUMOylation deregulates phosphorylation and ubiquitination in Alzheimer’s disease pathology. J Neurochem 2023;166(2):318–327

17. Ross RL, Corinaldesi C, Migneco G, Carr IM, Antanaviciute A, Wasson CW et.al, Targeting human plasmacytoid dendritic cells through BDCA2 prevents skin inflammation and fibrosis in a novel xenotransplant mouse model of scleroderma. Ann Rheum Dis 2021;80(7):920–929

18. Liu T, Zhang T, Nicolas M, Boussicault L, Rice H, Soldano A et al. The amyloid precursor protein is a conserved Wnt receptor. Elife 2021;10:e69199

19. Cui J, Wang X, Li X, Wang X, Zhang C, Li W et al. Targeting the γ/β-secretase interaction reduces β-amyloid generation and ameliorates Alzeimers’s disease-related pathogenesis. Cell Discov 2015;1:15021

20. Wasson CW, Abignano G, Hermes H, Malaab M, Ross RL, Jimenez SA, et al. Long non-coding RNA HOTAIR drives EZH2-dependent myofibroblast activation in systemic sclerosis through miRNA 34a-dependent activation of NOTCH. Ann Rheum Dis 2020;79:507–517

21. Hu X, Hou H, Bastain C, He W, Qiu S, Ge Y et al. BACE1 regulates the proliferation and cellular functions of schwann cells. Glia 2017;65(5):712–726

22. Niimi H, Pardali K, Vanlandewijck M, Heldin CH, Moustakas A. Notch signalling is necessary for epithelial growth arrest by TGF-beta. J Cell Biol 2007;176(5):695–707

23. Antinozzi C, Sgro P, Marampon F, Caporossi D, Del Galdo F, Dimauro I et al. Sildenafil counteracts the In Vitro activation of CXCL-9, CXCL-10 and CXCL11/CXCR3 axis induced by reactive oxygen species in scleroderma fibroblasts. Biology (Basel) 2021;10:491

24. Di Luigi L, Duranti G, Antonioni A, Sgro P, Ceci R, Crescioli C et al. The Phosphodiesterase type 5 inhibitor sildenafil improves DNA stability and redox homeostasis in systemic sclerosis fibroblasts exposed to reactive oxygen species. Antioxidants (Basel) 2020;9:786

25. Mouton-Liger F, Paquet C, Dumurgier J, Bouras C, Pradier L, Gray F et al. Oxidative stress increases BACE1 protein levels through activation of the PKR-eIF2α pathway. Biochim Biophys Acta 2012;1822:885–96

26. Hattori C, Asai M, Onishi H, Sasagawa N, Hashimoto Y, Saido T et al. BACE1 interacts with lipid raft protein. J Neurosci Res 2006;84(4):912–7

27. Liakouli V, Elies J, El-Sherbiny YM, Scarcia M, Grant G, Abignano G et al. Scleroderma fibroblasts suppress angiogenesis via TGF-β/Caveolin-1 dependent secretion of pigment epithelium-derived factor. Ann Rheum Dis 2018;77(3):431–440

28. Del Galdo F, Sotgia F, de Almeida CJ, Jasmin JF, Musick M, Lisanti MP et al. Decreased expression of caveolin 1 in patients with systemic sclerosis: crucial role in the pathogenesis of tissue fibrosis. Arthritis Rheum 2008;58(9):2854–65

29. Baranowski BJ, Hayward GC, Marko DM, Macpherson REK. Examination of BDNF treatment on BACE1 activity and acute exercise on Brain BDNF signalling. Front Cell Neurosci 2021;15:665867

30. Song Y, Wei J, Li R, Fu R, Han P et al. Tyrosine kinase receptor B attenuates liver fibrosis by inhibiting TGF-β/SMAD signalling. Hepatology 2023

31. George PM, Spagnolo P, Kreuter M, et al. Progressive fibrosing interstitial lung disease: clinical uncertainties, consensus recommendations, and research priorities. Lancet Respir Med. 2020;8(9):925–934

32. Khanna D, Clements PJ, Volkmann ER, et al. Minimal Clinically Important Differences for the Modified Rodnan Skin Score: Results from the Scleroderma Lung Studies (SLS-I and SLS-II). Arthritis Res Ther 2019;21(1):23

33. Watad A, Bragazzi NL, Tiosano S, Yavne Y, Comaneshter D, Cohen AD, et al. Alzheimer’s disease in systemic sclerosis patients: A nationwide population-based cohort study. J Alzheimer’s Dis 2018;65(1): 117–124

34. Ly PTT, Wu Y, Zou H, Wang R, Zhou W, Kinoshita A et al. Inhibition of GSK3β-mediated BACE1 expression reduces Alzheimer-associated phenotypes. J Clin Invest 2013;123:224–35

35. Estrach S, Ambler CA, Celso CL, Hozumi K, Watt FM. Jagged 1 is a beta-catenin target gene required for ectopic hair follicle formation in adult epidermis. Development 2006;133(22):4427–38

36. Dees C, Tomcik M, Zerr P, Akhmetshina A, Horn A, Palumbo K et al. Notch signalling regulates fibroblast activation and collagen release in systemic sclerosis. Ann Rheum Dis 2011;70:1304–10

37. Wasson CW, Ross RL, Morton R, Mankouri J, Del Galdo F. The Intracellular chloride channel 4 (CLIC4) activates systemic sclerosis fibroblasts. Rheumatology (Oxford) 2021;60(9):4395–4400

38. Munir H, Jones JO, Janowitz T, Hoffmann M, Euler M, Martins CP et al. Stromal-driven and amyloid β-dependent induction of neutrophil extracellular traps modulates tumour growth. Nat Commun 2021;12(1):683

39. Didier K, Giusti D, Le Jan S, Terryn C, Muller C, Pham BN et al. Neutrophil extracellular traps generation relates with the early stage and vascular complications in systemic sclerosis. J Clin Med 2020;9(7):2136

40. Cherubini E, Mariotta S, Scozzi D, Mancini R, Osman G, D’Ascanio M et al. BDNF/TrkB axis activation promotes epithelial-messenchymal transition in idiopathic pulmonary fibrosis. J Transl Med 2017;15:196

41. Huang J, Zhu H, Liu s, Li M, Li Y, Luo H, et al. Protein profiling in systemic sclerosis patients with different pulmonary complications using proteomic antibody microarray. Arthritis Res Ther 2024;26:29

42. LeRoy EC, Medsger TA Jr. Criteria for the classification of early systemic sclerosis. J Rheumatol 2001;28:1573–6

