## Supplementary Materials and Methods for "The Beta secretase BACE1 drives fibroblasts activation in Systemic Sclerosis through the APP/β-catenin/Notch signalling axis"

**Patient cell lines**

Full thickness skin biopsies were surgically obtained from the forearms of four adult healthy controls and four adult patients with recent onset SSc, defined as a disease duration of less than 18 months from the appearance of clinically detectable skin induration. All patients satisfied the 2013 ACR/EULAR criteria for the classification of SSc and had diffuse cutaneous clinical subset as defined by LeRoy et al (42). All participants provided written informed consent to participate in the study. Informed consent procedures were approved by NRES-011NE to FDG. Fibroblasts were isolated and established as previously described (2). Primary cells were immortalized using human telomerase reverse transcriptase (hTERT) to produce healthy control hTERT and SSc hTERT (2).

**Cell culture**

hTERT patient fibroblasts were maintained in Dulbecco’s modified Eagle medium (DMEM) (Gibco) supplemented with 10% FBS (Sigma) and penicillin-streptomycin (Sigma). Fibroblasts were treated with BACE1 inhibitors (M-3 (500nM-4μM, Sigma-Aldrich), AZD3839 (500nM-2μM, AstraZeneca) or XYT472B (10μM, company)) for 48 hours in a humidified incubator at 37C and 5% CO_2_. For the H_2_O_2_ the fibroblasts were treated with 100μM H_2_O_2_ for 24 hours. For BDNF stimulation, healthy and SSc dermal fibroblasts were grown in sera depleted media and stimulated with 20ng/ml BDNF for 48 hours.

**Morphogen stimulation**

Healthy dermal fibroblasts were serum starved for 24 h in DMEM containing 0.5% FBS and stimulated with 10 ng/ml TGF-β (R&D systems), 100 ng/ml Wnt-3a (R&D systems) or 100 nM SMO agonist (SAG) for 48 hr in combination with the BACE1 inhibitors.

**siRNA transfections**

A pool of four siRNAs specific for different regions of BACE1 or a negative control scrambled siRNA (Qiagen) were transfected into fibroblasts using Lipofectamine 2000 (Thermo Fisher). Fibroblasts were incubated for 48 h prior to harvesting.

**BACE1 overexpression**

Healthy dermal fibroblasts were transfected with 0.5µg of WT and Mutant BACE1 (Contained within pcDNA3.1 mammalian expression vector) (7) using lipofectamine 2000. Fibroblasts were incubated for 48 h prior to harvesting.

**Caveolin-1 shRNA**

Caveolin-1 was silenced by shRNAmir GIPZ lentiviruses transduction (Open Biosystems, Surrey, UK). Briefly Healthy dermal fibroblasts were transduced with lentiviruses containing scramble control shRNA and Caveolin-1 shRNA and incubated for 48 hours. Transduced fibroblasts were selected with 1.0μg/ml puromycin (Life Technologies) for 10 days (28).

**Gel Contraction Assay**

200,000 fibroblasts were suspended in Collagen Gel Working Solution (Cambridge Biosiences, Cat. CBA-201). After collagen polymerization, 1ml of 1%FBS culture medium was added on top of each collagen gel and incubated overnight at 37C with 5% CO2. Next day, the sides of the cell-collagen gels were detached using sterile spatulas ensuring that the gels were circular. Images and weight of the gels were collected after 72 hours.

**Western blotting**

Total proteins were extracted from fibroblasts in RIPA buffer and resolved by SDS-PAGE (10-15% Tris-Glycine). Proteins were transferred onto Hybond nitrocellulose membranes (Amersham biosciences) and probed with antibodies specific for alpha smooth muscle actin (Abcam), BACE1 (Sigma), Notch 1 (Cell signalling), GLI2 (R&D systems), β-catenin (cell signalling), phosho-SMAD3 (S423/S425) (Abcam), Caveolin 1 (Santa-Cruz), Jagged 1 (Cell signalling) and β-Actin (Sigma). Immunoblots were visualized with species-specific HRP conjugated secondary antibodies (Sigma) and ECL (Thermo/Pierce) on a Biorad chemiDoc imaging system.

**Quantitative Real time PCR**

RNA was extracted from cells using commercial RNA extraction kits (Zymo Research). RNA (1ug) was reverse transcribed using cDNA synthesis kits (Thermo). QRT-PCRs were performed using SyBr Green PCR kits on a Thermocycler with primers specific for BACE1 (Forward: GCAGGGCTACTACGTGGAGA, Reverse: GTATCCACCAGGATGTTGAGC), alpha SMA (Forward; TGTATGTGGCTATCCAGGCG Reverse; AGAGTCCAGCACGATGCCAG), Collagen type 1A2 (Forward; GATGTTGAACTTGTTGCTGAGC Reverse; TCTTTCCCCATTCATTTGTCTT), CTGF (Forward; GTGTGCACTGCCAAAGATGGT Reverse; TTGGAAGGACTCACCGCT), Notch 1 (Forward; CCAGAACTGTGAGGAAAATATCG Reverse; TCTTGCAGTTGTTTCCTGGAC), Hes1 (Forward; TACCCAGCCAGTGTCAAC Reverse; CAGATGCTGTCTTTGGTTTATCC), GLI2 (Forward; TTTATGGGCATCCTCTCTGG Reverse; TTTTGCATTCCTTCCTGTCC) and GAPDH (Forward; ACCCACTCCTCCACCTTTGA Reverse; CTGTTGCTGTAGCCAAATTCGT). Data were analysed using the ΔΔ Ct method. GAPDH served as a housekeeping gene.

**Immunohistochemistry**

Immunohistochemistry was performed as previously described (17). Sections were stained with a BACE1 antibody (1/200) (Sigma), visualised using an HRP conjugated rabbit secondary and counterstained with haematoxylin. An anti-Rabbit Isotype control antibody was used as a negative control and this staining was performed alongside the BACE1 staining.

**Bleomycin Mouse Model**

Mice used in the bleomycin study were severe combined immunodeficient (CB17/Icr-Prkdcscid/IcrIcoCrl, Charles River). 100µl of bleomycin (200µg/ml in PBS) was injected into a single location on the shaved back of the mice. Bleomycin was administered once every other day for 3 weeks. The mice were euthanized and the skin was harvested using a punch biopsy (16).

**BACE1 KO mouse study**

Mice were given free access to food and water and maintained on a 12 h light/dark cycle (PPL PP2103311). BACE1 KO mice, C57BL/6J background, were obtained from the Jackson Laboratory (B6.129-Bace1tm1Pcw/J) and BACE1 KO and wild type control mice were generated using a BACE1 heterozygote breeding strategy (9). Punch biopsies (3mm) were obtained from the skin of these mice and fibroblasts were isolated from the biopsies, as previously described (17).

**Lentiviral Transduction**

Fibroblasts (FBs) were grown from healthy control forearm biopsies and immortalised using retrovirus expressing human telomerase (hTERT) as previously outlined (2). HOTAIR expression was then induced by transduction with GIPZ lentiviruses carrying HOTAIR gene sequence or scrambled RNA sequence as control in frame with puromycin resistance gene and GFP fluorochrome gene (Open Biosystems, Surrey, UK). For this purpose FBs were seeded at 50% confluence and infected with lentiviral particles in serum free DMEM and incubated for 6h, after which an additional 1ml of DMEM containing 10% FCS was added and the cells were incubated for a further 72h. Stably transduced FBs were positively sorted for GFP fluorescence employing Fluorescence Activated Cell Sorting in sterile conditions (BD INFLUX). Positively sorted cells were further selected in media containing 1.0μg/ml puromycin (Life Technologies) for 10 days.

**Patient clinical characterization and serum sample collection**

Consecutive SSc patients who were enrolled at the University of Leeds between January 2012 and December 2014 were included in this study and longitudinally evaluated every six months up to 10 years. Two control groups included respectively HCs and patients with sRP defined for anti-centromere or anti-Scl70 positivity without any other manifestations of connective tissue disease.

The medical history collection, a comprehensive clinical examination, the annual pulmonary function tests and echocardiogram, were used to define clinical characteristics, including disease cutaneous variant and duration from the first non-Raynaud symptom, anti-centromere and anti-Scl70 antibody positivity, severity of skin sclerosis according to modified Rodnan skin score (mRSS), visceral organ involvement. High-resolution computed tomography and right-heart catheterization were performed when interstitial lung disease (ILD) or pulmonary hypertension presence or progression were clinically suspected. Twenty-four-month ILD progression according to Erice working group criteria, 12-month mRSS progression according to minimal clinically important difference (MCID) definition in patients with diffuse cutaneous variant (dcSSc), and 10-year SSc-related mortality, were selected as outcome measures.

**Luminex Analysis of Patient Sera**

BDNF, AB40 and AB42 serum protein analysis performed by the CLIA certified Myriad Rules-Based Medicine (Austin, TX, USA) using a Multi-Analyte Profiling (MAP) multiplexed immune Luminex assay.

**Statistical analysis**

Categorical variables were reported as numbers and percentages while continuous variables as mean±SD, mean ±standard error (SE), or median with interquartile range (IQR). Comparisons of continuous variables were performed with Student’s t-test or Wilcoxon rank-sum test, as appropriate, with Benjamini-Hochberg adjustment in case of multiple comparisons. The Kaplan-Meier method with log-rank test was conducted to determine if there were differences in the survival distributions according to pre-set clinical risk categories. Statistical significance was defined as a *p*<0.05 for all the statistical analyses. All the tests were two-tailed. Data were analysed using RStudio (version 2022.02.3+492).

**Ethics Approval and consent to participate**

All participants provided written informed consent to participate in this study. Informed consent procedure was approved by NRES-011NE to FDG by the University of Leeds
